# HIV-1 infection promotes neuroinflammation and neuron pathogenesis in novel microglia-containing cerebral organoids

**DOI:** 10.1101/2024.06.13.598579

**Authors:** Srinivas D. Narasipura, Janet P. Zayas, Michelle K. Ash, Anjelica Reyes, Tanner Shull, Stephanie Gambut, Jeffrey R. Schneider, Ramon Lorenzo-Redondo, Lena Al-Harthi, João I. Mamede

**Affiliations:** Department of Microbial Pathogens and Immunity, Rush University Medical Center, Chicago, Illinois, USA; Department of Medicine, Division of Infectious Diseases, Northwestern University Feinberg School of Medicine, Chicago, Illinois, USA; Center for Pathogen Genomics and Microbial Evolution, Institute for Global Health, Northwestern University Feinberg School of Medicine, Chicago, Illinois, USA

**Keywords:** Cerebral organoids, microglia, HIV, neuropathogenesis, neuroinflammation, iPSC, HPC

## Abstract

Cerebral organoids (COs) are a valuable tool to study the intricate interplay between glial cells and neurons in brain development and disease, including HIV-associated neuroinflammation. We developed a novel approach to generate microglia containing COs (CO-iMs) by co-culturing hematopoietic progenitors and induced pluripotent stem cells. This approach allowed for the differentiation of microglia within the organoids concomitantly to the neuronal progenitors. CO- iMs exhibited higher efficiency in generation of CD45^+^/CD11b^+^/Iba-1^+^ microglia cells compared to conventional COs with physiologically relevant proportion of microglia (∼7%). CO-iMs exhibited substantially higher expression of microglial homeostatic and sensome markers as well as markers for the complement cascade. CO-iMs showed susceptibility to HIV infection resulting in a significant increase in several pro-inflammatory cytokines/chemokines and compromised neuronal function, which were abrogated by addition of antiretrovirals. Thus, CO-iM is a robust model to decipher neuropathogenesis, neurological disorders, and viral infections of brain cells in a 3D culture system.

## Introduction

Neurodegenerative diseases are debilitating disorders characterized by deterioration of brain function and affect millions of people worldwide. Currently, the most-utilized models to study brain diseases involve *postmortem* human brain tissue^1,2^, non-human primates (NHP)^3^, small animal models^4–7^, *in vitro* 2-dimentional (2D) cultures of primary cells (mostly fetal in origin)^8^ or cell lines^9–11^. While relevant in the context of their specific studies, many of these models are limited by availability, cost, and the differences between species,^5,9^. Cerebral organoids have emerged as a tool to study the human brain in health and disease ^10,11^. COs provide a three-dimensional (3D) architecture of resident brain cells (neurons, astrocytes, oligodendrocytes, and microglia), albeit the lack vascularization of the tissue environment. They are developed from embryonic stem cells (ESCs) or induced pluripotent stem cells (iPSCs) to form 3D tissue organization and spatial arrangement of diverse CNS cell types that recapitulate the intricate pattern and functionality of the brain tissue^12–15^. Organoids of various parts of the brain, such as forebrain, midbrain, and hippocampus, are a powerful tool to study neurodegenerative diseases, neurodevelopmental disorders, and the interplay between brain cells under homeostatic and diseased conditions^16^. As the genetic manipulation of iPSCs is very well established, COs allow for the study of specific molecular mechanisms related to brain biology and neurovirology. Furthermore, COs express genes associated with human cortical development, and the developing neurons display active neural networks^17,18^.

Despite their progress as a powerful model platform to study neuronal diseases at an organ level, COs lack proper abundance of cell types that are of non-ectodermal origin, such as microglia^19^. To circumvent this, short term co-cultures of pre-differentiated microglia are engrafted to organoids; however, the extent to which these incorporated microglia survive and function in these structures is unknown^20–23^.

Human immunodeficiency virus (HIV) invades the brain as early as 8 days post-infection^24–26^, initiating an inflammatory cascade leading to neuronal injury clinically manifesting as HIV associated neurocognitive disorders (HAND). Close to 50% of people living with HIV (PLWH) suffer from HAND^27,28^, a spectrum of neurocognitive impairments that range from mild cognitive and motor impairments to severe dementia. HAND can significantly affect the quality of life and daily functioning of those impacted^29^. While mechanisms driving HAND are multifaceted, low level of HIV in the brain under antiretroviral therapy (ART) contributes to persistent neuroinflammation^29,30^. Evidence of pockets of replication in peripheral tissues despite suppressive ART exist, where the presence of low levels of viral RNA, DNA, and proteins, is thought to be the primary driver of neuroinflammation^31^.

There are numerous models to study HIV neuropathogenesis, each with its own limitations. While human samples are ideal for studying HIV in the brain, post-mortem samples of the brain are often difficult to obtain, and these cohorts are usually heterogeneous in terms of their treatment regimens and comorbidities, as well as the method of preservation utilized during brain harvest. It is also difficult to conduct longitudinal assessment of brain tissue from PLWH. Non-human primates have served as an important model of HIV/SIV neuropathogenesis, yet SIVmac and SHIVs are not genetically identical to HIV-1 and there are profound differences in species-specific responses including differential molecular aspects to viral replication, disease progression and responses^32,33^. Humanized mouse models while having human immune cells, contain vast majority of host cells in the brain that are resistant to HIV infection^34^. These studies are also conducted in the backdrop of graft versus host disease (GVHD)^35^, which develops in all humanized models and has to be controlled for^34,36^. These caveats to existing models to study HIV in the brain underscore the need to develop models of brain cells in 3D culture systems that recapitulate the human condition.

Microglia, the resident myeloid cells of the brain, are the major cell type infected by HIV in the CNS^37,38^. Astrocytes, the most abundant glial cells in the brain, are infected to a lesser extent and have mechanisms to suppress HIV transcription^39–42^. Macrophages and infiltrating T cells are also known to harbor HIV in the brain^43–45^. Development of appropriate models to dissect the role of microglia in HIV infection becomes paramount not only to understand HIV associated neuroinflammation, but also to study the brain as a reservoir for HIV. This brain reservoir contributes to the overall reservoir of HIV in the body, as HIV can egress from the brain to seed other lymphoid tissues^46^. Recently, Tang *et al*., have shown that microglia isolated from adult human brain are susceptible to HIV infection and lack significant viral genetic diversity, indicating the microglial reservoir is seeded early in infection^47^. However, these studies exploit monolayer cell culture of microglia in the absence of other cell types. Using innately developed microglia in a CO model, Ormel *et al*. demonstrated HIV-1 infection of microglia^48^. In this model, the limitations include presence of lower numbers of microglia (≤1%) compared to the human brain and hence requirement of a high amounts of virus (100ng/CO) to achieve a successful infection. dos Reis *et al*., engrafted infected microglia (adult primary or a cell line) into COs^49^. While this method is adequate for incorporating microglia into COs, it has several drawbacks, as the microglia engrafted into COs are previously infected rather than infected after incorporation into the tissue environment, and the study uses cell cultures that are difficult to obtain for large-scale experiments. Recently, in an elegant study, Schafer *et al*., xenotransplanted erythromyeloid progenitors incorporated COs into adult NOD/SCID mouse brain and showed generation of functionally mature human microglia that can operate within a physiologically relevant vascularized brain environment^50^. Although this model can provide excellent insights into the biology and pathology of human microglia, the laborious surgery methods make it impractical to conduct large scale studies. Further, complications to study HIV pathogenesis may arise due to resistance of mouse immune system to HIV infection and fast occurrence of GVDH to humanization.

Hematopoietic progenitors (HPCs) also known as erythromyeloid progenitors (EMPs) derived from iPSC differentiate into microglia when properly cued^51^. Utilizing this principle, we developed an approach to generate COs containing microglia by combining HPCs with iPSCs during the first stage of embryoid body formation. We demonstrate successful proliferation, differentiation, and maturation of incorporated HPCs into microglia in mature COs. These cerebral organoids containing microglia (CO-iMs) express high levels of microglia specific homeostatic and sensome markers. They were successfully infected with HIV and exhibited augmented neuroinflammatory characteristics with infection. Further, by incorporating constitutively expressing red fluorescent protein (RFP) into HPCs, we demonstrate that microglia in CO-iMs are specifically amenable to genetic modifications.

## Materials and Methods

### Culture and maintenance of iPSCs

Human iPSC lines were obtained from Rutgers University Cell and DNA Repository (RUCDR) Infinite Biologics (ID# NN0003920, source; male, fibroblast) and Coriell Institute for Medical Research (Camden, NJ; ID#AG27602, Source; male, fibroblast) maintained in complete mTeSR+ medium (Stem Cell Technologies; Vancouver, Canada) containing 0.1% penicillin/streptomycin and propagated as small aggregates using accutase or ReLeSR (Stem Cell Technologies). All iPSC colonies were cultured at 37°C in a 5% CO2 incubator on 6-well tissue culture plates coated with matrigel (Corning, NY). Coating of plates was performed by thoroughly mixing matrigel thawed on ice with ice cold DMEM/F-12 medium at 1:100 dilution followed by incubating 1ml/well for 1h at room temperature (RT) or 37°C in 5% CO2. iPSCs were routinely validated for markers, OCT4, TRA-1-60 and Ki-67 using flow cytometry.

### Labelling of iPSC with turboRFP and generation of HPCs

TurboRFP expressing lentiviral particles under the control of EF1a promoter (EF1A-RFP-LV) was purchased from Cellomics (Halethorpe, MD, # PLV-10072). iPSC from RUCDR (termed iPSC- Ru) were dissociated into single cells with accutase and plated at 0.3x10^6^ cells per well of a 6-well plate in mTeSR+. Next day, cells were replaced with fresh mTeSR+ and infected with EF1A-RFP- LV viral particles at 0.1 to 1.0 transduction units/cell. Three days later, wells were replaced with fresh mTeSR+ media supplemented with puromycin (0.75 μg/ml). Media was changed every other day and healthy colonies formed under lowest viral concentrations were expanded for at least two more passages under puromycin (6-8 days per passage) before freezing the cells for stocking or differentiating into HPCs.

Both wild type (WT) and RFP labeled iPSCs were differentiated to HPCs using STEMdiff hematopoietic kit (Stem Cell Technologies). Briefly, iPSCs were dissociated into small colony clumps via ReLeSR treatment (incubating as per STEMCELL protocol, followed by minimal gentle pipetting) and plated at different dilutions of colonies per well of a 6-well plate in mTeSR+. On the next 24 to 48h after confirming for optimal size and number of colonies (5-10 small colonies/cm^2^), mTSR+ media was replaced with fresh media A and the protocol was followed for the next 12-14 days as per kit instructions (Stem Cell Technologies). On day 12 and day 14, round floating HPCs were harvested and cultured for up to three more days in X-vivo-15 basal media (Lonza, Switzerland), supplemented with 100 ng/mL M-CSF (Invitrogen, Massachusetts), 25 ng/mL IL-3 (R&D), 2 mM glutamax (Invitrogen), 0.055 mM β-mercaptoethanol (Invitrogen), 100 U/mL penicillin, and 100 μg/mL streptomycin. HPSCs phenotype was confirmed by flow cytometry before the cells were used for experiments or frozen for later usage. HPCs were routinely validated for expression of markers; CD45, CD43, and CD34 via flow cytometry.

Images of iPSC colonies and HPCs were captured under bright field or fluorescence using a Keyence BZ-810 fluorescence microscope (Keyence, IL, USA) at 20x magnification. Cells were frozen in Cryo-SFM freezing media (Promo Cell, Heidelberg, Germany).

### Generation of cerebral organoids

Cerebral organoids were generated using STEMdiff cerebral organoid kit (Stem Cell Technologies) with minor modifications^15,45^. On day 0, iPSC colonies were suspended into single cells using accutase. To generate a regular CO, 10,000 iPSCs were incubated in organoid formation media; alternatively, to generate a CO-iM, 10,000 iPSCs were co-cultured with 3,000 HPCs and incubated in organoid formation media. Ultra-low binding, round bottom 96-well plates (Corning) were used for embryoid body (EB) formation. Heat stable recombinant human bFGF (Gibco) at 5U/ml was supplemented to organoid formation media only on day 0. In the case of CO-iMs, macrophage colony stimulating factor (MCSF, 50 ng/ml) and interleukin 3 (IL-3, 20 U/ml) were used throughout organoid development to support survival, proliferation and differentiation of HPCs to microglia. In order to prevent the fusion of multiple organoids during the maturation stage (around day 20), each organoid was separated into single wells in low binding 24-well plates with approximately 700 μl maturation media per organoid and incubated at 37°C, 5% CO2 on an orbital shaker at 90 rpm/min. Anti-adherent solution (Stem Cell Technologies) was used to create low binding plates.

### Single cell dissociation of organoids and flow cytometry

Organoids were gently washed with sterile HBSS, cut into small pieces, and incubated with papain (30 units/ml, Stem Cell Technologies) along with human recombinant DNAseI (20 U/ml, Roche) for 20-30 min at 37°C, 5% CO2 on an orbital shaker at 90 rpm/min. Preparation and activation of papain solution was performed as per the instructions (Stem Cell Technologies). One ml of papain solution per organoid per well in a 12-well plate was used. Approximately, every 10 minutes, organoids were dissociated by vigorous trituration (10∼15 times) using a 1 mL pipette tip. By 30 min, most of the organoids dissociated into single cells and the remaining small tissue and cell aggregates were gently pressed using the back of a 1ml syringe, and triturated. The whole cell suspension was passed through a cell strainer (40 μM) and washed with 2-3ml of sterile HBSS. The flow through containing single cells were centrifuged at 1300rpm for 8 min, washed again with sterile HBSS, resuspended in 1ml HBSS, counted using a cell counter and processed for flow cytometry. This protocol consistently resulted in >80% of cell viability as measured by tryphan blue or calcein violet-AM (BioLegend).

iPSC colonies cultured on matrigel coated plates were detached with accutase and washed twice with sterile PBS. Floating HPCs were also washed twice with sterile PBS. Single cells from iPSC, HPCs or organoids were resuspended in 100 μl of PBS, incubated with Fc block (Cat #564220, BD Biosciences, NJ) for 10 min at RT as per instructions and subjected to either cell surface staining for surface markers using staining buffer (BD Biosciences), or intracellular staining using perm/fix reagents (BD Biosciences). For intracellular staining, cells were incubated with antibodies at 4°C for 60 min in the dark, washed three times thoroughly with perm/wash buffer, resuspended in perm/wash buffer and run on a Fortessa flow cytometer (BD Biosciences). Primary conjugated antibodies used for flow are listed in supplementary Table 1. Data was analyzed using FlowJo software (v10.0.0) and normalized to the respective isotype IgGs or fluorescent minus one (FMOs) controls. For compensation, 2 drops of anti-mouse Ig, k beads (BD Biosciences) were incubated with primary antibodies for 20 min at 4°C, washed once with PBS, then a drop of negative control BD comp beads (BD Biosciences) was added at the end and fixed with 2% paraformaldehyde. FlowJo was used to set compensation parameters.

### RNA and DNA extraction from organoids

Organoids were stored in RNAlater (500 μl/organoid) at −20°C until processing. RNA extraction was performed using the RNeasy mini kit (Qiagen, Hilden, Germany). Briefly, organoid was quickly brought to room temperature, RNA later was carefully removed and replaced with 500 μl of RLT lysis buffer in sterile tubes containing lysing matrix D beads (Cat #6913100, MP Biomedicals). Tubes were placed in bead beater (BeadBlaster 24, Bechmark Scientific, NJ) at a speed of 4 M/S, for 30 sec with 15 sec intervals for 5 cycles to completely disintegrate and dissolve organoids in RLT buffer. Further steps were followed according to kit protocol. Column DNAse digestion was performed, and elution was done using only 40 μl of elution buffer. RNA was quantified using a nanodrop and samples that had 260/280 ratio of >1.8 and 260/230 ratios between 2.0-2.2 were selected for further processing. Complementary DNA (cDNA) was synthesized using a Qscript kit (QuantaBio, MA) as per kit instructions in a T-100 (Bio-Rad) thermocycler.

For DNA extraction, organoids were removed from RNAlater, cut into small pieces and suspended in RLT buffer (700 μl/organoid) containing proteinase K. Samples were incubated at 56°C for 30 min-2h with intermittent vortexing until all the pieces were dissolved. Next steps were followed according to DNAeasy Kit (Qiagen). DNA was eluted in 40 μl elution buffer and quantified using a nanodrop. Samples with 260/280 ratio of >1.8 and 260/230 ratios between 2.0-2.2 were considered for further processing.

### Quantitative real time PCR

Real time quantitative RT-PCR (qRT-PCR) was performed on cDNA samples (at least 1/20 volume) using SYBR green master mix. Samples were loaded on a 96-well fast block and run on a QuantStudio 5 Flex (Thermo Fisher) instrument. Protocol settings were as follows: hold stage (95 °C for 10 minutes), 45 cycles of PCR stage (95 °C for 20 seconds and 60 °C for 1 minute) and melt curve stage (95°C for 15 seconds, 60 °C for 1 minute and 95 °C for 15 seconds). Primers used are listed in supplementary Table 2. Cycle threshold (Ct) values were subtracted with Ct values of housekeeping genes (GAPDH and 18S), relative expressions of genes were computed using 2^-ΔΔCt^ method and are expressed as fold changes compared to respective control groups.

Similarly, real time quantitative PCR (qPCR) was performed on DNA samples using Taqman based primers and probes for HIV DNA and human DNA essentially as previously described^52^. Ct values of HIV amplifications were normalized to human DNA and are expressed as fold changes compared to respective control groups.

### HIV infection of cerebral organoids

Infectious particles of HIV-1Ba-L (Cat# ARP-510, BEI Resources) were produced in peripheral blood mononuclear cells (PBMCs) isolated from a healthy seronegative donor using Ficol- Hypaque density gradient centrifugation. Approximately 100 million cells were cultured in complete RPMI 1640 medium which included 10% heat-inactivated fetal bovine serum (Gemini Bio Products, Calabasas, CA), 2 mM L-glutamine (GlutaMAX supplement; ThermoFisher), 20 U/mL IL-2 (AIDS Reagent Program, Germantown, MD) and 1% penicillin/streptomycin (Sigma-Aldrich, St. Louis, MO), Cells were activated for 3 days with 1 μg/mL/million cells of purified NA/LE mouse anti-human CD3 and CD28 (BD Biosciences) and infected with HIVBaL at 0.01 ng/10^6^ cells. PBMCs were monitored to express at least 6–10% intracellular p24^+^ in T lymphocytes before harvesting the supernatant. Supernatant was spun at 5000 rpm for 10 min to remove cell debris and concentrated via Lenti-X as per the instructions (Takara Bio). HIV-Gag-iGFP_JRFL (the following reagent was obtained through the NIH HIV Reagent Program, Division of AIDS, NIAID, NIH: Human Immunodeficiency Virus (HIV) Gag-iGFP_JRFL, ARP-12456, contributed by Dr. Benjamin Chen), containing an eGFP detailed in fused to GAG was produced and purified as previously ^53,54^. Purified viruses were quantified using a HIV1 p24 ELISA kit (#ab218268, Abcam, Waltham, MA). Approximately, 60 to70-day old organoids were infected with 10ng/ml concentration of HIV in a low-binding 24 well plates at one organoid/well/700 μl of maturation media containing MCSF and IL-3. Three days later, the wells were gently washed and replaced with fresh maturation media. cART treatment consisted of plasma levels of nevirapine (NVP, 10 µM) and dolutegravir (DTG, 3.90 μg/mL) that were added 24 h prior to infection, at the time of infection, and after day 3 wash.

### Immunofluorescence of organoids

Organoids were flash frozen fresh in cryomolds containing Tissue-Tek OCT (optimal cutting temperature) compound. Organoid specimens were washed twice in PBS and acclimated to OCT for few minutes in a small weight boat. Then, organoids were transferred into a labeled cryomold where they were oriented and fully immersed in OCT while avoiding bubble formation. The cryomolds containing the OCT immersed organoids were immediately transferred and submerged for about 10 seconds into a cooling bath consisting of a beaker containing 2-propanol inside a dry ice bucket until the OCT compound solidified or was visibly frozen. OCT blocks were kept on dry ice until stored at −80°C. or sectioned (10-20µm) into cover slips using a microtome cryostat and stored at −80°C.

Sections were subject to multiplex immunofluorescence (mIF) staining for target cells and imaged by widefield deconvolution or spinning disk confocal microscopy.^55^ mIF consisted of a series of consecutive rounds of staining and bleaching after initial fixing and blocking of tissues. Briefly, sections in cover slips were mounted and glued into home-made polyurethane resin staining chambers and fixed for 5 minutes at room temperature in the dark using a PIPES/PFA fixing solution (PIPES buffer pH 6.8, and 10% paraformaldehyde). After removing the fixing solution, tissues were washed three times with PBS and blocked for 1 hour at room temperature using donkey serum blocking buffer (PBS, donkey serum, 10% sodium azide, and 10% triton X- 100). All staining cocktails were prepared in blocking buffer and sections were stained for 1 hour at room temperature or overnight at 4°C. Additionally, bleaching was accomplished by exposing the tissues to light for 1 hour at room temperature in bleaching buffer (7 volumes of distilled water, 2 volumes of 1M NaHCO3, and 1 volume of 30% H2O2). After removing the bleaching solution, tissues were washed three times with PBS and kept on PBS at 4°C, or immediately used for a new round of staining. Antibodies used to stain COs are indicated and listed in supplementary Table 3. Ashlar (v1.18.0 https://github.com/labsyspharm/ashlar) was used to align max intensity projection of the Z-stacks (0.5um spacing) for the multiple mIF rounds with max pixel deviation of 200 and filter sigma of 2.

Images were acquired with a Nikon wide-field Ti-e2 equipped with an LED light source (Spectra X) and a PCO edge 4.2BI camera using standard Chroma DAPI/GFP/TRITC/Cy5 polychroic and emission filters for DAPI/GFP/TRITC with oil 40x magnification or with a Nikon Ti-e2 spinning disk confocal (Crest X-Light V3) equipped with a celesta laser light source. Widefield images obtained with the widefield Nikon TIe-2 microscope were deconvolved with FlowDec using the Lucy-Richardson algorithm with the support of Pims relying on python 3.x using calculated PSFs from FlowDec as previously reported ^56,57^.

### Multiplex ELISA

Luminex assay for simultaneous quantification of multiple cytokines and chemokines (custom built, premixed kit) in the organoid supernatants were performed in a 96-well microplate format as per kit instructions (Luminex performance assay, R & D Systems). Data capture was done using a Luminex Flexmap 3D analyzer (ThermoFisher) and data analysis used a five-parameter logistic (5-PL) curve fit of a standard curve per analyte.

### Bulk RNA sequencing (RNAseq), Single cell RNA sequencing (scRNAseq), and bioinformatics analysis

Total RNA was extracted from organoids using the RNeasy kit (Qiagen) and checked for concentration, RIN/RQN and 28S/18S ratios using Agilent 4150 TapeStation system (Agilent Technologies). Transcriptome sequencing was performed by DNBSEQ™ sequencing technology platform (BGI/Ionomics) which includes stranded library preparation, 100 bp paired-end sequencing and ≥30million reads. Sequencing data was demultiplexed and trimmed using Trimmomatic v0.36 to remove adapters and low-quality reads. Trimmed reads were aligned to the Homo sapiens reference genome GRCh38 and transcripts quantified using the Hisat2-StringTie pipeline^58^. Differential gene expression analysis of the quantified gene transcripts was performed with DESeq2 v.1.42.0 R package using R v.4.3.2. After retaining genes with nonzero total read count, we identified differentially expressed genes (DEGs) either between CO-iMs and COs; HIV- infected and uninfected CO-iMs; or between ART-treated and untreated HIV-infected CO-iMs using as cut-offs an absolute log2 fold change > 1 and a false discovery rate (FDR) < 0.05 using the Benjamin-Hochberg procedure. Gene enrichment analyses for each comparison were subsequently performed using gene set enrichment analysis (GSEA) to identify specific Gene Ontologies (GO), KEGG, and REACTOME pathways associated with CO-iMs and/or HIV infection and ART treatment. We performed GSEA using clusterProfiler v.4.10.0 in R with all lists of genes ranked by the corresponding log2 fold change. For these analyses all genes whose gene symbols could be mapped to ENTREZ Ids using the org.Hs.eg.db v.3.18.0 Bioconductor annotation package were included.

For scRNAseq four CO-iMs were individually processed for isolation of single cells using papain and DNAseI treatment aspreviously described. The single cell suspensions from the organoids were resuspended in Parse cell buffer containing BSA, manually counted using a hemocytometer and assessed for cell viability by calcein violet-AM. The samples were centrifuged at 1300 rpm for 10 minutes and then fixed/permeabilized using the Evercode fixation v2 kit (Parse Biosciences, ECF2101) and stored at −80°C according to the instructions. The bar-coding and library preparation for single cells were performed using the spit-pool based approach (Evercode™ WT Mini v2, Parse Biosciences). In brief, ∼27,000 fixed cells from each organoid were pooled and underwent three rounds of barcoding. The final pooled cells were divided into 2 equal libraries for cDNA synthesis and library construction to recover 10,000 cells. Multiplexed libraries were sequenced using an Illumina NovaSeq X Plus system, with a 10B Flowcell and PE150 sequencing. Fastq files were generated from bcl files and processed using the ‘Parse Biosciences analysis pipeline’ using the GRCh38 reference genome to generate a gene expression matrix. After quality filtering, 6,669 and 6,331 estimated cells for each library were analyzed with 92,691 and 91,357 reads per cell, respectively. 40,666 and 40,588 genes were analyzed with a median of 4,387 and 4,279 genes per cell for each library. Seurat v.5.0.3 R package was used for analysis. We filtered cell barcodes with reads from less than 200 genes or more than 7,500, as well as barcodes with more than 10% of mitochondrial genes that resulted in 11,440 remaining barcodes for analysis. We used the harmony method with 5000 variable features and 30 principal components to integrate the dataset from the two libraries and the Leiden method for clustering with 0.3 resolution using 10 principal components. DEGs that were upregulated in each cluster and expressed by a minimum of 25% of the cells in that cluster compared to the rest of the cells were identified using Seurat’s implementation of the MAST algorithm Finally, ScType method R function using the curated ScType database filtered for brain tissue excluding cancer cellswas used to annotate the clusters obtained.

### Statistics

Statistical analyses were performed using Prism software (GraphPad Prism, San Diego, CA). Comparisons between groups were done using an unpaired, non-parametric, Mann-Whitney test. Experiments were conducted independently at least three times, and the data represented as standard error of mean (SEM). p value ≤ 0.05 is considered statistically significant. * Represents p≤ 0.05, ** represents p≤ 0.01 and *** represents p≤ 0.001.

## Results

### Microglia are efficiently differentiated in organoids produced by combining iPSC with HPCs during EB formation

Given that HPCs (or erythromyeloid progenitor cells, EMPs), when provided with proper cues can differentiate into microglia *in vitro*^58^, we hypothesized that by mixing HPCs with iPSCs during EB formation stage and providing appropriate conditions (MCSF and IL-3) allows HPCs to proliferate and differentiate into microglia in line with organoid maturation. We first confirmed iPSC lines to be of high quality by morphology (colonies with tight borders with less than 5% differentiation, **Suppl. Fig. 1A**) and by phenotype (expression of classical markers, OCT-4 and TRA-1-60 pluripotency markers, and Ki-67 proliferation marker, **Suppl. Fig. 1B**). We then differentiated unlabeled, or RFP labeled, iPSCs into HPCs and confirmed for its small spherical floating cells morphology (**Suppl. Fig. 1C**), and expression of key progenitor marker, CD43 (>80% of cells) along with stemness marker, CD34 (∼60% of cells) and myeloid marker, CD45 (**Suppl. Figs. 1D and E**). In addition, efficient labeling of iPSCs and HPCs with constitutively expressing turboRFP was also confirmed (**Suppl. Figs. 1F-I**). Next, we generated unguided cerebral organoids (COs), as described^12,48^ and generated CO-iMs by mixing iPSCs with HPCs at a 10:3 ratio on day 0 and incubated with MCSF (50ng/ml) and IL-3 (20U/ml) throughout the formation, expansion and maturation stages. The schematic in **Figure 1A** outlines both approaches. We did not detect noticeable changes in the morphological features of organoids at any stage by adding HPCs in CO-iMs from the conventional COs developed only from iPSCs (**Fig. 1B**). Further, incorporation of HPCs did not affect the size of the COs at any measured developmental stage (**Fig. 1C**). Cyclic multiplex immunofluorescence (mIF) imaging of approximately 60 to 70-day old CO-iMs revealed robust presence of TMEM119^+^ as well as Iba1^+^ microglia cells (**Figs. 1D-F and Suppl. Figs. 2A-D**). Furthermore, mIF confirmed the presence of similar numbers of astrocytes (GFAP^+^/S100β^+^) and neurons (Tuj1^+/^NeuN^+^) in CO-iMs and COs. Additional lineage specific markers for glial and neuronal cells were used as well, which revealed the presence of differential marker expression on these cell types, for example not all GFAP^+^ are S100β^+^. Similarly, not all NeuN^+^ are Tuj1^+^ or MAP2^+^ (**Figs. 1D-F and Suppl. Figs. 2A-D**). We quantified astrocytic (GFAP^+^ and S100β^+^), neuronal (Tuj1^+^) or NPC (Nestin^+^) populations in these organoids via flow cytometric analysis (**Suppl. Figs. 2E-H**). Incorporation of HPCs did not affect the relative proportion of NPC (∼30-37% Nestin^+^), astrocytes (∼38-40% S100b^+^ and ∼45-55% GFAP^+^), or neuronal cells (∼45-50% Tuj1^+^) between COs and CO-iMs (**Suppl. Figs. 2I and J**). However, there was a significant increase (∼4.6 fold) in myeloid (CD45^+^) cells in CO-iMs compared to COs (**Figs. 2A and B**). In line with these findings, we observed a significant increase (∼3.5-4 folds) in CD11b/Iba1 double positive cells (**Figs. 2C**). As expected, CD11b/Iba1 double positive cells were only found in CD45^+^ population and they were devoid in CD45^-^ population (**Fig. 2A**, bottom dot plots). Interestingly, within the CD45^+^ myeloid population, there were similar proportions of CD11b/Iba1 single positive or double positive cells, both in COs and CO-iMs (**Fig. 2D**). In parallel, in organoids generated using iPSCs-CRL, there was a significantly higher (∼2 folds) proportion of CD11b/Iba1 double positive cells; however, there was no difference in CD45^+^ population in CO-iMs when compared to COs (**Figs. 2E and F**). In addition, in CO-iMs generated with incorporated HPCs stably expressing RFP, most of RFP^+^ cells were CD45^+^ (>60%, **Figs. 2G and H**). As previously documented^44^, we found CD45^+^/Iba1^+^ microglia population in COs, albeit to lower levels (**Figs. 2B, C, F and** H). Taken together, these results demonstrate that HPCs incorporated at the beginning of EB formation will successfully proliferate and differentiate into mature microglia at the same timeframe with the overall growth and maturation of the neurons and astrocytes in the brain organoids.

**Figure 1:**
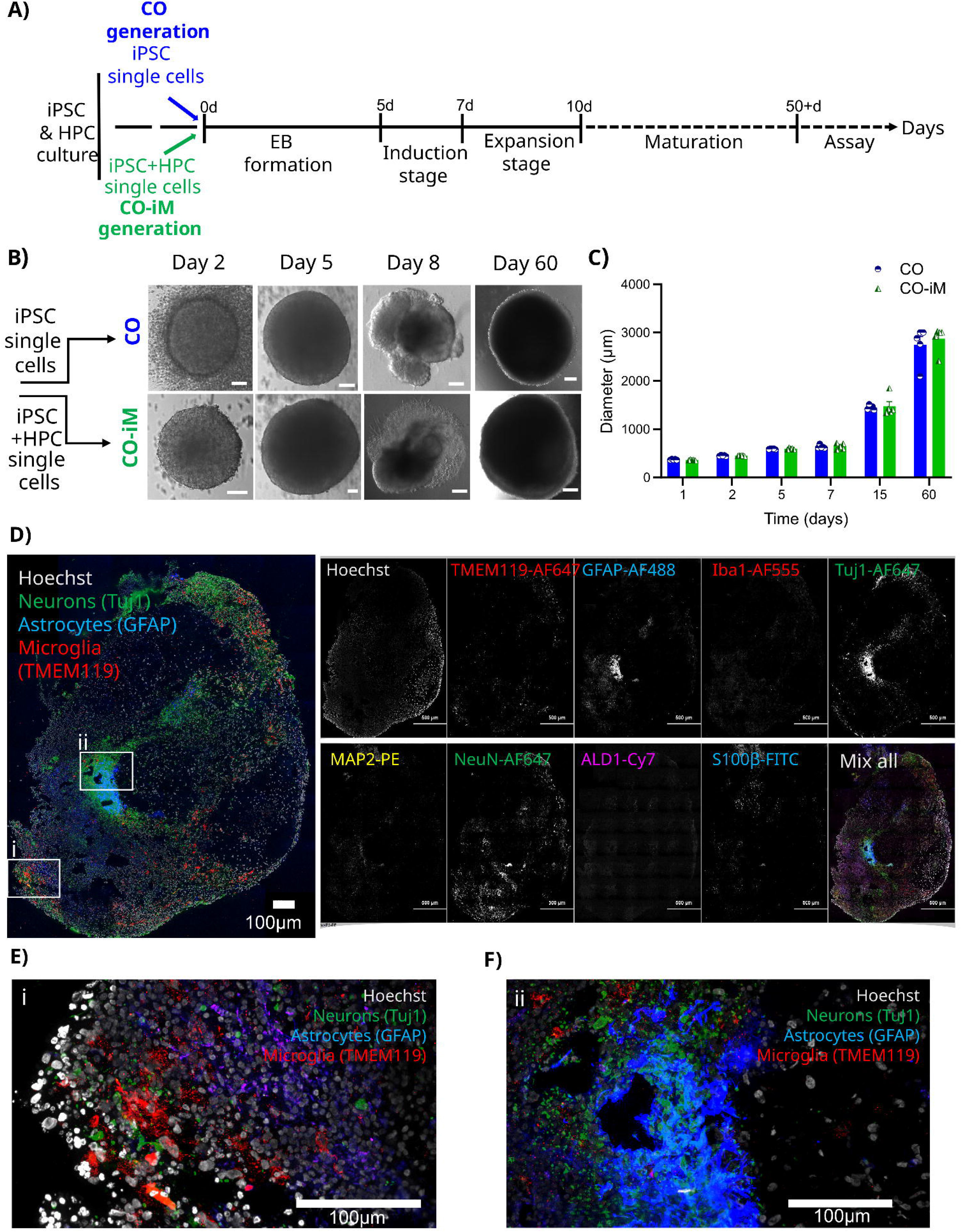
Generation and characterization of CO-iMs. **(A)** Schematic flow diagram indicating the timeline as well as all developmental stages for generation of CO and CO-iMs. **(B)** Representative bright field images of a CO and a CO-iM through different stages of their development. Scale bars, 100 μm. **(C)** Diameter sizes of CO and CO-iMs measured at key time points during their differentiation (n=5 per time point). **(D)** Left, representative multiplex IF image of a CO-iM stained for nucleus (Hoechst, white), microglia (TMEM119, red), neurons (green, Tuj1) and astrocytes (GFAP, blue) Scale bar, 150 μm. Panel on the right showing individual staining for indicated markers and a merged image from all. Scale bars, 100 μm. **(E and F)** Zoomed insets of i), and ii) regions in (D). Scale bars, 100 μm.

**Figure 2:**
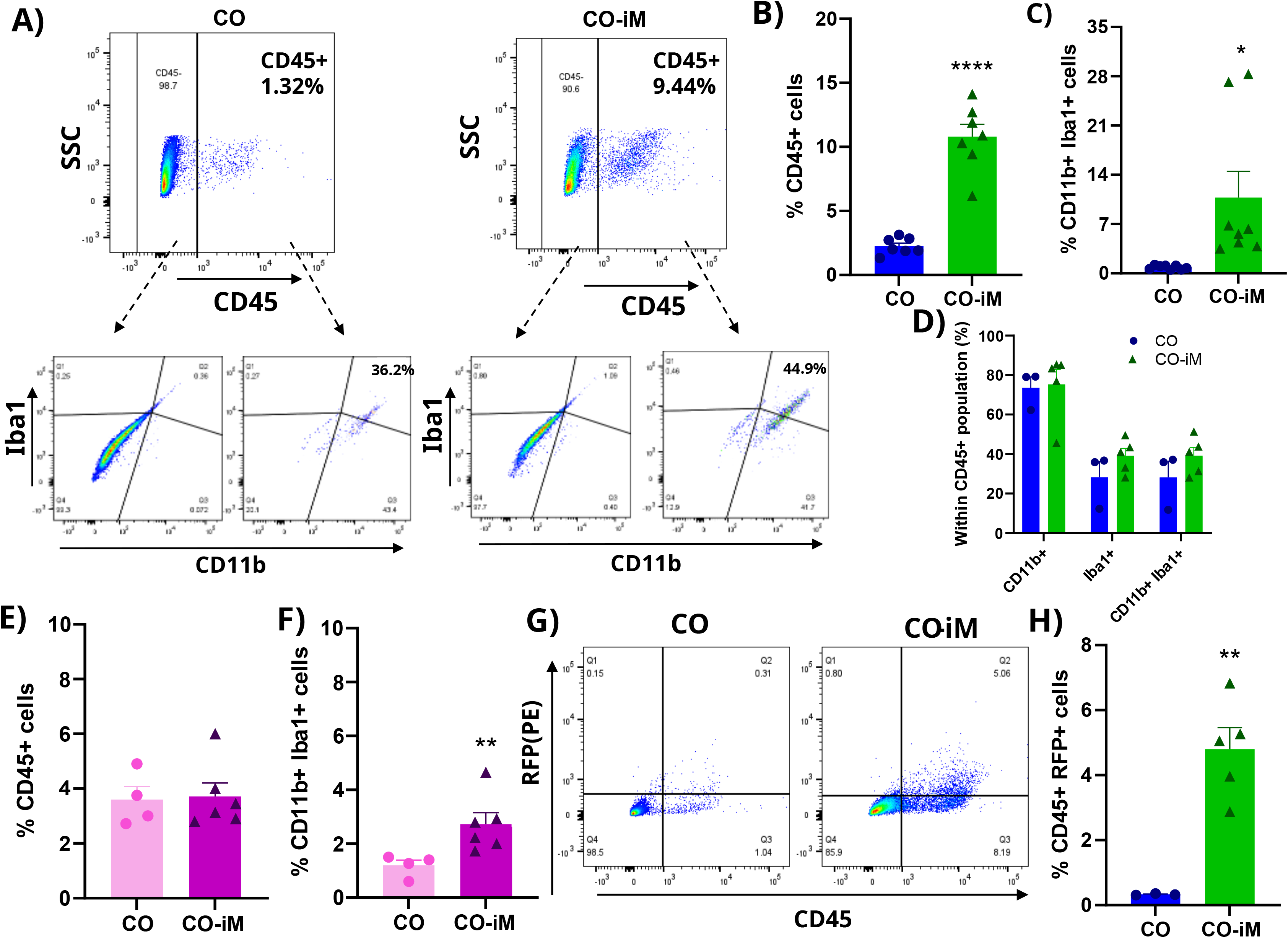
Phenotypic characterization reflecting enrichment of microglia in CO-iMs. CO and CO-iMs were processed and dissociated into single cell suspension, stained, and analyzed for cell surface and intracellular proteins by flow cytometry. **(A)** Representative dot plots showing CD45^+^ and CD45^-^ populations from total cells (top), and CD11b^+^ and Iba1^+^ populations (bottom). Arrows indicate populations analyzed within CD45^+^ or CD45^-^ gates. **(B and C**) Cumulative data indicating percent CD45^+^ and CD11b /Iba1 double positive cells in total population of organoids generated using iPSC-Ru line (n=7 for COs and CO-iMs, respectively) **(D)** Cumulative data showing percent CD11b/Iba1 single positive as well as double positive cells within CD45^+^ population in organoids generated using iPSC-Ru line (n=3 for COs, and n=5 for CO-iMs). (**E and F)** Cumulative data showing percent CD11b/Iba1 single as well as double positive cells within CD45^+^ population in organoids generated using iPSC-CRL line (n=4 for COs, and n=6 for CO-iMs). **(G)** Representative dot plots showing that majority of RFP^+^ cells are also CD45^+^ cells. **(H)** Cumulative data indicating percent RFP^+^ CD45^+^ population (n=3 for COs, and n=5 for CO-iMs). Each symbol represents an individual organoid. Data are presented as mean ± SD using non-parametric Mann-Whitney test. *p < 0.05, **p < 0.01, ***p < 0.001.

### Transcription analysis exhibit induced expression of key microglia specific genes in CO-iMs

To further characterize the myeloid cells in CO-iMs for proliferation, abundance, differentiation and maturation, we quantified transcriptional profiles of important homeostatic microglia markers such as *CD11B*, *IBA1* and *TMEM119*^59^ along with *CD45*, a myeloid marker expressed at lower levels compared to monocytes/macrophages in steady state microglia^60^. We found that expression levels of all these specific genes were significantly higher in CO-iMs when compared to conventional COs. The trend was similar when normalized to either *GAPDH* (**Fig. 3A**) or *18S* endogenous controls (**Suppl. Fig. 3A**). Interestingly, in CO-iMs, the level of *CD45* induction was much lower compared to *CD11B* or *IBA1* (∼8-10 fold vs 60-100 fold; **Fig. 3A**), suggesting that low expression of *CD45* in combination with high *CD11B* perhaps indicates a resting steady-state feature of microglia in CO-iMs^60,61^. Expression levels of astrocyte markers (*S100β, ACSBG1* and *EAAT1*), neuronal progenitor/immature neuron markers (*Vimentin* and *Nestin*) and mature neuronal markers (*MAP2* and *NEUN*) were similar between CO-iMs and COs (**Figs. 3B-D** and **Suppl. Fig. 3B).** Further, *APOE*, a lipid transporter, mainly expressed in astrocytes was also found to be unchanged (**Fig. 3B** and **Suppl. Fig. 3B).**

**Figure 3:**
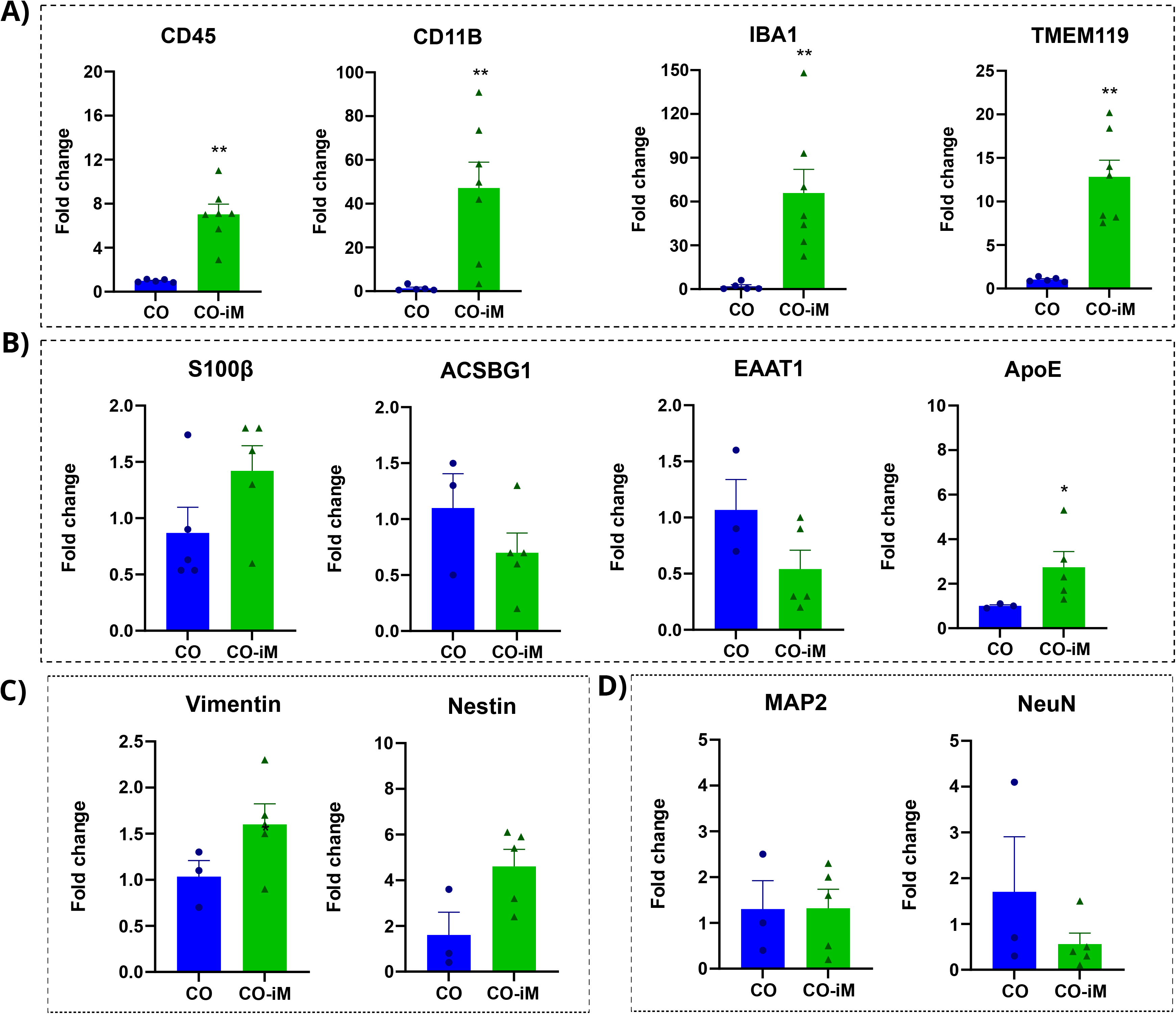
Genotypic characterization reflecting enrichment of microglia markers in CO-iMs. qRT-PCR analysis of COs (n=3-5) and CO-iMs (n=5-7). mRNA levels were normalized to the endogenous reference GAPDH and expressed as fold change relative to CO using 2^-(ΔΔCt)^. Fold change for each sample in COs was assessed by subtracting its delta Ct from average delta Ct, followed by 2^-(ΔΔCt)^. **(A)** mRNA levels of homeostatic microglia markers: *CD45*, *CD11B*, *IBA1* and *TMEM119*. **(B)** mRNA levels of astrocyte markers: *S100b*, *ACSBG1*, *EAAT1* and *APOE.* **(C and D)** mRNA levels of neuronal progenitor and mature neuronal markers: *Vimentin, Nestin*, *MAP2* and *NeuN*. Each symbol represents an individual organoid. Data are presented as mean ± SD using non-parametric Mann-Whitney test. *p < 0.05, **p < 0.01, ***p < 0.001.

Similar observations with respect to gene expressions were noted in organoids generated using iPSC-CRL line. Wherein, expression of *CD11B* and *IBA1* were significantly higher in CO- iMs in comparison to COs (**Suppl. Figs. 3C and D**). *S100b* and *Nestin* levels were unchanged between CO and CO-iMs (**Suppl. Figs. 3E and F**). The expression levels of *CD45* mRNA and *TMEM119* mRNAs were not significantly different between COs and CO-iMs with this iPSC line (**Suppl. Figs. 3C and D**) as it was found with CD45 protein expression (**Fig. 2E**). These findings reflect inherent intra- and inter-group variations as observed among batches and to the different iPSCs cell lines^14,18,62^.

To assess the differences between COs and CO-iMs, we characterized the global transcriptome profile and compared COs and CO-iMs by RNAseq analysis. Gene enrichment analysis using differentially expressed genes demonstrated striking differences in genes involved in immunity such as phagocytosis, leukocyte migration, chemotaxis, cell migration, and others (**Fig. 4A**) and pathways such as complement and coagulation, interactions of cytokine receptors with cytokines and foreign antigens, TLR signaling, etc. (**Fig. 4B**). Differential gene expression (DGE) analysis identified 1412 (5.6% of the total number of genes tested) genes that showed significant upregulation in CO-iMs compared to COs (FDR<0.05 and log2 fold-change>1) and 1486 (5.9% of the total tested) downregulated genes, indicating significant changes associated to the inclusion of microglia in the COs (**Fig. 4C and Suppl. File 1**). Subsequently, we performed a Gene Set Enrichment Analysis (GSEA) with the entire list of quantified genes ranked by log2 fold change of the CO-iM vs CO comparison. We observed a significant enrichment of genes related to complement, cytokine expression, and innate immunity (**Figs. 4A** and **B**) amongst genes upregulated in CO-iMs when compared with COs (**Fig. 4C**). While enrichment for genes related to neuronal and astrocytic homeostatic function was detected amongst the downregulated genes. Interestingly, we find genes with important brain functions to be statistically significantly upregulated in CO-iMs, such as *CDA* and *PKD2L1* that have BBB and oligodendrocyte myelination functions, respectively. Conversely, *LINGO4*, a gene anticipated to operate either upstream or within the positive regulation of synapse assembly, is downregulated, which might point to important microglia molecular function in synapse regulation as reviewed^64^.

**Figure 4:**
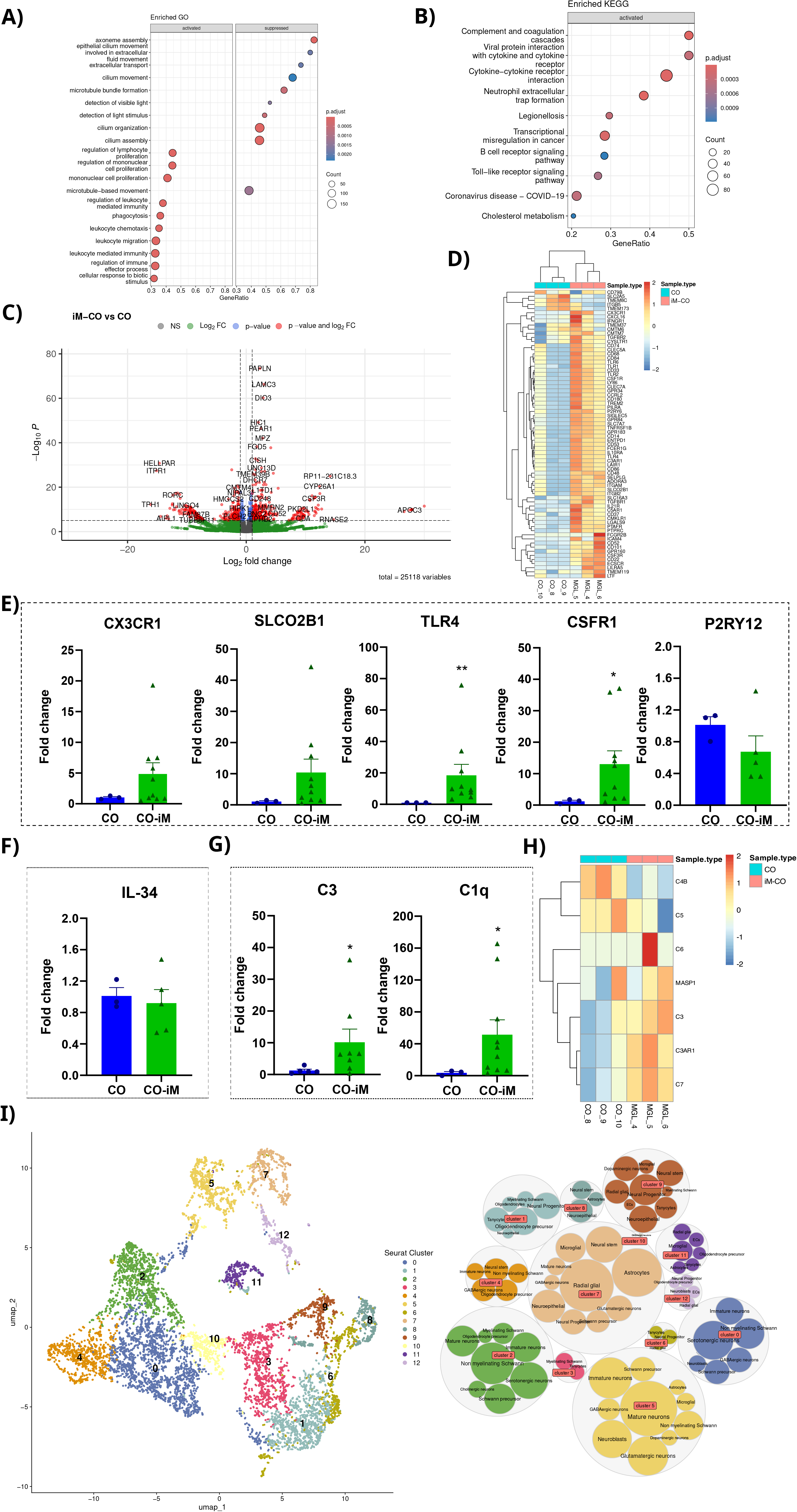
Transcriptional Profile of CO and CO-iMs. RNA-seq analysis of ∼65-day old iPSC- Ru line derived COs (n = 3) and CO-iMs (n = 3). **(A)** Dot plot showing the enriched GO terms of biological processes and molecular functions amongst activated and suppressed genes identified using absolute log2 fold change > 1 and a false discovery rate (FDR) < 0.05 using the Benjamin-Hochberg procedure in CO-iMs when compared to COs. **(B)** Dot plot showing the enriched REACTOME pathways (FDR <0.1) in activated as well as suppressed genes in CO-iMs when compared to COs. For (**A**) and (**B**) the x-axis indicates the ratio of genes identified by our analysis and the total number of genes that constitute the pathway, the size of the dot is based on gene count in the pathway, and the color of the dot shows the pathway enrichment significance. **(C)** Volcano plot comparison of CO-iMs vs COs indicating average log2 (fold change) versus log10 (FDR) for all genes. Genes upregulated and downregulated by 2-fold change and FDR < 0.05 are labeled with red dots. **(D)** A heatmap showing expression of microglia sensome markers in individual replicates of CO-iMs and COs, (n = 3 for CO-iMs and n = 3 for COs). Color indicates the expression level scaled for each gene by centering and scaling. **(E)** qRT-PCR quantification to determine mRNA levels of microglia immune sensome markers: *CX3CR1*, *SLCO2B1*, *TLR4*, *CSF1-R* and *P2RY12* **(F)** qRT-PCR quantification to determine mRNA levels of *IL34*, an essential cytokine for microglia maturation and survival. **(G)** qRT-PCR quantification to determine mRNA levels of *C3* and *C1Q*, components of complement pathway. **(H)** Heatmap showing expression of microglia complement pathway genes in individual replicates of CO-iMs and COs, (n = 3 for CO- iMs and n = 3 for COs). Color indicates the expression level scaled for each gene. **(I)** Uniform Manifold Approximation and Projection (UMAP) plot for dimension reduction of 11,440 barcoded single cells derived from 4 individual CO-iMs (∼120 day old) subjected to scRNA seq analysis exhibiting 12 distinct clusters as identified by graph-based clustering of cell-specific gene expression. Subsequent cell annotation allowed us to define the most likely cell type corresponding to each cluster. The bubble plot shows all the cell types that were considered by ScType for cluster annotation that were assigned to at least 100 cells with the exception of cluster 10 where no assignment reached that cut-off and the only assignment obtained is shown. The outer bubbles correspond to each cluster with size reflecting the number of cells in the cluster, while the inner bubbles correspond to considered cell types for each cluster, with the biggest bubble corresponding to assigned cell type, except for cluster 3 and 10 where we did not obtain high quality assignments.

Microglia sense changes in their surroundings for immune regulation and host defense functions. Therefore, we quantified the expression of microglia-specific sensome genes listed from a well-established cluster^65,66^. Heatmap analysis indicated significant enrichment for the majority of these genes in CO-iMs when compared to COs (**Fig. 4D**). Validation by qRT-PCR analysis for some of these genes including *CX3CR1*, *TLR4*, *SLCO2B1*, *P2RY12, CSF1-R* and *TMEM119* exhibited significant induction of all these genes in CO-iMs compared to COs, except for *P2RY12* (**Figs. 3A, 4E** and **Suppl. Figs. 3A and 4A**). MCSF and IL34 are required for the survival, proliferation, and maturation of microglia^61,62^. CSF1-R is the common receptor for MCSF and IL34. We provided MCSF along with IL-3 throughout the culture period. Substantial induction of *CSF1- R* mRNA in CO-iMs (**Fig. 4E** and **Supp. Fig. 4A**) along with similar yet noticeable levels of *IL34* mRNA in both types of organoids (**Fig. 4F** and **Suppl. Fig. 4B**) indicated a self-sufficient milieu, not only for initial proliferation and differentiation of HPCs into microglia, but also for survival, proliferation, and maturation of microglia in our 60+ day old, cultured organoids. Microglia plays an important role in synapse pruning (removal of excess and less active synapses), via classical complement pathway, in a developing brain. Synapse pruning is pivotal in maintenance and refinement of synapses^67^. We assessed the expression levels of *C1Q* and *C3*, two early key players of complement pathways via qPCR. These two mRNAs were significantly elevated in CO-iMs (**Fig. 4G** and **Suppl. Fig. 4C**). Similar observation was made in CO-iMs generated from iPSC- CRL, except for *C1Q,* as it did not change between two types of organoids (**Suppl. Figs. 4D**). In addition to *C3*, heatmap analysis indicated induction of additional members of complement pathways including *C7*, *C3AR1* and *MASP1* in CO-iMs (**Fig. 4H**). While *C6* was unchanged between CO and CO-iMs, *C5* and *C4B* were significantly downregulated in CO-iMs (**Fig. 4H**), suggesting a homeostatic maintenance of complement cascade. Collectively, these results demonstrate that microglia generated in this manner in CO-iMs exhibit adult-like features including expression of homeostatic and immune surveillance genes.

While targeted approaches and bulk RNAseq capture the overall trends of the CO-iMs, and mIF can capture the different expression profiles and different spatial cellular distributions, they are less powerful in detecting unknown or less abundant cell types and subtypes. Thus, to capture the molecular identity of different cell types more accurately, we isolated single cells from four <u>C</u>O-iMs (approximately 4-month-old), pooled into 2 groups and performed scRNAseq on a total of 13,000 cells. Graph-based cell clustering based on transcriptional patterns detected 13 distinct clusters that we could assign to 11 main CNS cell types (**Fig. 4I**), as indicated by the bigger bubbles within each cluster in Fig. 4I, including various types of neurons (mature, serotonergic and others), microglia (**Suppl. Fig. 3F**), astrocytes (**Suppl. Fig. 3H**), oligodendrocyte precursor cells, and others. A heat map with the most highly expressed genes in each cluster is shown (**Suppl. Fig. 3G**). Taken together, scRNAseq confirmed the emergence of mature microglia in CO-iMs, assuring that CO-iMs presented here are a good representation of different brain cells fortifying the importance of these cell culture 3D models to neural and neurovirology research.

### Microglia are robustly infected with HIV in CO-iMs

As CO-iMs contain sufficient and physiologically relevant proportions of microglia cells, we used CO-iMs as a model to study HIV infection. Brain organoids have been previously used to model HIV, CMV and ZIKV infections^48,49,63,64^. However, for HIV infections, mature microglia were infected first and then allowed to traffic into COs^48,49^. Doing so will not allow us to study the complete establishment, spread, and consequences of HIV infection at its different stages. To address the questions, on the levels of microglia susceptibility to HIV and the neuropathology within an organoid to closely mimic natural mode of infection in the CNS, CO-iMs were infected with two different infectious HIV isolates, HIV-1Ba-L (R5 tropic) and HIV-Gag-iGFP_JRFL (neurotropic envelope where the viral particles are green due to fusion of a fluid phase GFP within Gag)^65^. HIV infection did not affect the morphology of the organoids up to the measured 30 days post-infection (**Figs. 5A and B**), as reported for other viral infections^64,66,67^. Extracellular p24 quantification by ELISA indicated a significantly higher amount of virus released from CO-iMs when compared to COs. This release was inhibited in the presence of cART, which was also used as a control for viral input in the supernatant (**Fig. 5C**). Similarly, qPCR on these infected organoids showed a 6 to 8-fold increase in HIV DNA when compared to COs (**Fig. 5D**). A representative amplification plot for viral and human DNA is shown in **Figure 5E**. These data demonstrate a robust productive infection in CO-iMs in comparison to COs. Cyclic mIF analysis demonstrated co-localization of HIV p24^+^ (mAb staining or iGFP-Gag+) in Iba^+^ microglia cells (**Figs. 5F-I**). Furthermore, we observed viral particles in regions surrounding microglia infected cells and regions with heavy concentration of viral particles surrounding microglia that are not infected (arrows with dotted lines **Fig. 5H**). Flow cytometry-based quantification analysis indicated that HIV-iGFP+ (or p24^+^) cells were positive for CD45^+^, while the CD45^-^ population was negative (**Figs. 5J and K**). We also report significantly higher CD45^+^/p24^+^ cells in CO-iMs compared to COs (**Fig. 5L**). Further, by flow cytometry we found that the majority of HIV^+^ (iGFP+) cells were CD11b^+^ which was almost undetectable after ART treatment (**Suppl. Fig. 5A**). Our findings demonstrate robust HIV infection in microglia, which are also the major source of productive infection in CO-iMs fully modeling results reported from post-mortem brains and humanized mice studies.

**Figure 5:**
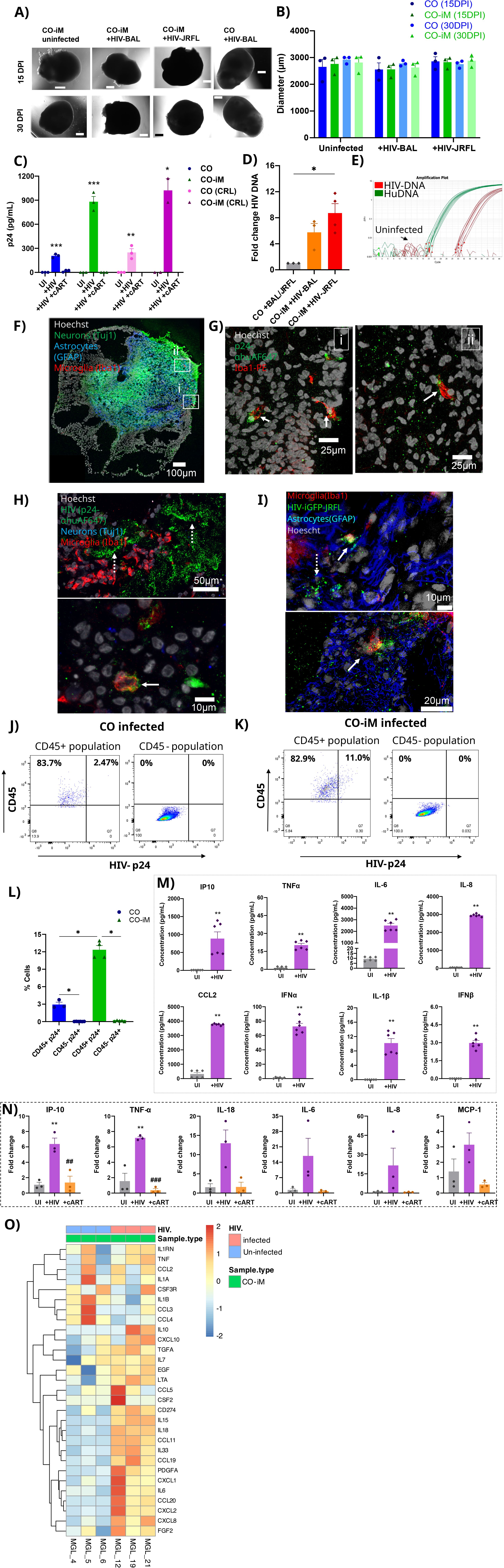
Robust response of microglia to HIV-1 infection in CO-iMs. **(A)** Representative bright field images showing the morphology of CO and CO-iMs (∼60 day old) after 15 or 30 days post infection (DPI) with HIV-1 or mock. Scale bars, 500 μm. **(B)** Diameter sizes of CO and CO- iMs measured 15 DPI or 30 DPI with HIV or mock (n=2 for COs, and n=3 for CO-iMs). **(C)** Organoids were infected with HIVBaL (at 10 ng/ml) on day 0. At 3 DPI, organoids were washed once with PBS and replaced with fresh media. Then, at 5 DPI, the culture supernatant was collected and used to quantify p24 by ELISA. n=3 for each condition with organoids generated from two different hiPSC cell lines (RUCDR and CRL, respectively). **(D)** Organoids were infected with HIVBaL or HIV-iGFP-JRFL as explained in (C), genomic DNA was extracted and quantified for HIV DNA using Taqman qPCR and represented as fold change to COs infected with HIVBaL or HIV-iGFP_JRFL (n=3 for COs, n=3 for CO-iM + HIVBaL, and n=4 for CO-iMs+HIV-iGFP_JRFL). **(E)** A representative Taqman qPCR amplification plot showing amplification of human DNA as well as HIV-GAG for all samples shown in (D). **(F)** Representative mFI images of a CO-iM showing IBA1^+^/HIV^+^ microglia after 5 DPI HIV infection. Scale bars, 100 μm. **(G)** Zoomed in insets of regions i) and ii) from (G) showing p24^+^ (green), Iba1^+^ (red) cells. Scale bars, 25 μm. **(H** and **I)** mFI images of infected CO-iMs exhibiting p24^+^ (green) and Iba1^+^ (red) co-expressing cells (arrows with solid lines) as well as free virus (arrows with dotted lines) nearby Iba1^+^ cells. Scale bars, 50 and 10 μm, respectively. **(J and K)** Organoids were infected with HIVBaL for 5 days as explained, dissociated into single cell suspension and stained for cell surface (CD45) and intracellular (p24) markers for flow cytometry analysis. Representative dot plots for CD45 and p24 populations in COs (J, left) and in CO-iMs (K, left). **(L)** Cumulative data indicating percentage of p24^+^ cells in CD45^+^ and CD45^-^ populations, respectively. n=3 for COs, and n=4 for CO-iMs. **(M)** Multiplex ELISA to quantify various pro-inflammatory cytokines/chemokines in the supernatants of CO-iMs infected with HIVBaL for 5 days (n=6), or mock (n=6) for 5 days. **(N)** qPCR analysis for mRNA expression of various pro-inflammatory cytokines/chemokines (as indicated in the figures) in CO-iMs either infected with HIVBaL for 5 days, or mock infected, or infected and treated with cART. n=3 in each category. (O) Heatmap showing relative expression of a list of inflammatory cytokines/chemokines in infected vs uninfected CO-iMs (n = 3 for each group). Color indicates the expression level scaled for each gene. Each symbol represents an individual organoid. Data are presented as mean ± SD using non- parametric Mann-Whitney test. *p < 0.05, **p < 0.01, ***p < 0.001.

### HIV infection of CO-iMs results in heightened inflammatory condition

To determine the immunological and cellular responses from HIV infection, we examined the expression of inflammatory cytokines at the RNA and protein levels by bulk RNA sequencing, qRT-PCR, and multiplex ELISA. Multiplex ELISA indicated significantly higher secretion of several inflammatory cytokines such as IL-6, IP-10, TNFα, IL-8, MCP1, INFα, IL1β and INFβ in the supernatant of CO-iMs infected with HIV compared to non-infected CO-iMs (**Fig. 5M**). We then evaluated the mRNA expression of these cytokines under similar conditions via qRT-PCR.

We found that many of these cytokine transcripts were induced in response to HIV infection, including *IP10* and *TNFα*, which were statistically significant (**Fig. 5N and Suppl. Fig. 5B**). In parallel, heatmap analysis via RNAseq, showed a strong induction of many of the cytokines and chemokines in CO-iMs infected with HIV when compared to non-infected controls, including IL6, IP10 and TNFα (**Fig. 5O**), which was abrogated by cART treatment (**Fig. 5N and Suppl. Figs. 5B, C**). While *IL1β, INFβ, and IL1α* mRNAs showed no relative increase at the measured timeframes post HIV infection (**Fig. 5N** and **Suppl. Fig. 5D)**; their measured protein levels significantly increased (**Fig. 5M**), which may reflect a longstanding permanence of these cytokines in this system after transcriptional shutdown.

Transcriptome analysis between HIV infected and non-infected CO-iMs showed strong upregulation and activation of immune responses (IFNs signaling), *IL-10* (log2FC = 5.65), *IL-6* (log2FC = 3.58), *IL-33* (log2FC = 2.834) and antigen processing and presentation (**Figs. 6A and B**). Conversely, HIV-1 infection resulted in suppression of pathways of glucuronidation, neuron fate commitment, neuropeptide functions, axon and forebrain development, and lipoprotein function (**Figs. 6A and B**). Employing DEG analysis, we identified a total of 1826 upregulated genes (7% of the total number of genes tested) (FDR<0.05 and log2 fold-change>1) and 2158 (8.3% of the total tested) downregulated genes with HIV infection, indicating significant responses to infection in CO-iMs (**Fig. 6C and Suppl. File 2**). Of note, we find that caspase-5, a protein that is associated with pyroptosis, cell death, and the activation of the inflammasome as a response to neuronal trauma resulting in cell death^68–70^ is highly upregulated with infection (log2FC = 9.26). Further, DTHD1, a widely uncharacterized protein found to be involved in apoptotic events in the brain and found to be active and highly expressed in the brains of Alzhiemer’s and Multiple Sclerosis lesions^71,72^ is highly upregulated (log2FC = 23.83). In addition, other transcripts that are important for neuronal homeostasis and function, such as psynaptophysin (log2FC = −1.7); *MAP2* (log2FC = -1.77), and *EAAT2* (log2FC = -1.46) were downregulated approximately between 60- 70% with infection. Strikingly, *NOS1* (log2FC = 22.34), a gene that is involved in nitric oxide synthesis from L-arginine, which has many neurotransmitter properties and has been shown to relate to neurotoxicity and neurodegenerative diseases was also highly upregulated with HIV infection (**Fig. 6C and Suppl. File 2**). Interestingly, cART treatment with and without infection only mildly reduced the expression of DEGs (<1%, **Suppl. Fig. 5E and Suppl. File 3**).

**Figure 6:**
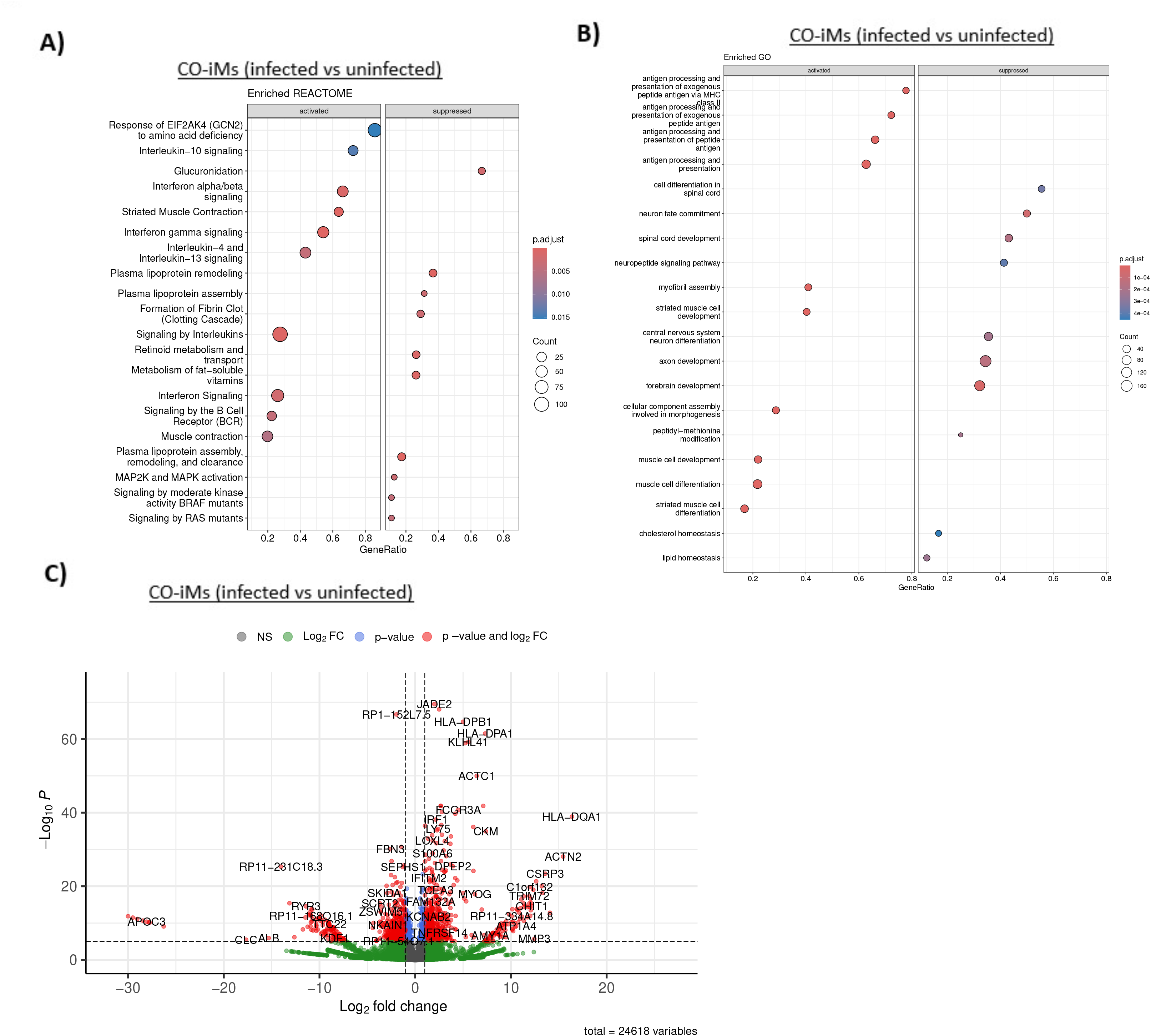
Transcriptional Profile of CO and CO-iMs infected with HIV. RNA-seq analysis of 65-day old iPSC-Ru line derived CO-iMs infected with HIVBaL (n = 3) for 5 days or mock infected (n = 3) or infected+cART treated (n = 3) for 5 days. **(A)** REACTOME dot plot showing the pathways (FDR <0.1) enriched in activated and suppressed gene in infected CO-iMs when compared with their uninfected counterparts. **(B)** Dot plot showing the GO terms (FDR <0.05) of biological processes enriched in activated and suppressed genes in infected CO-iMs when compared with their uninfected counterparts. For (A) and (B) the x-axis indicates the ratio of genes identified by our analysis and the total number of genes that constitute the pathway, the size of the dot is based on gene count in the pathway, and the color of the dot shows the pathway enrichment significance.  **(C)** Volcano plot comparison of infected vs uninfected CO-iMs indicating average log2 (fold change) versus log10 (FDR) for all genes. Genes upregulated and downregulated by 2-fold change and FDR < 0.05 are labeled with red dots.

Taken together, these data demonstrate robust infection and replication of HIV in CO-iMs, driven primarily by HIV infection of microglia in this system. The infection drives inflammatory responses in these organoids that lead to poor neuronal health. Furthermore, we find that HIV-1 infection and replication of CO-iMs have an impact in neuronal function and development, modeling one of the hallmarks of HIV-1 infection, where there is a big biological impact in the brain.

## Discussion

We developed a microglia containing CO model by co-culturing iPSCs and HPCs from the same line, at the beginning stage of EB formation. In this 3D culture model, HPCs proliferate and mature into microglia, as indicated by robust expression of microglia specific homeostatic markers, as well as sensome markers in 60+ day old organoids. In the same system and in parallel, iPSCs differentiate into NPCs, which further differentiate into mature neurons and astrocytes. Notably, by introducing HPCs at the formation stage along the 3D formation stage (expansion to maturation stage), rather than the distinct engraftment of fully differentiated microglia we gain unique advantages such as: (i) mimicking the brain cell composition and cellular interactions from the formation of CO-iMs; (ii) use of less HPCs/microglia per organoid as this method does not depend on the efficiency of microglia post-engraftment; and, (iii) the ability to regulate the number of HPCs and in turn microglia, per organoid. Further, by harnessing the self-organization capability of iPSCs instead of NPCs we generated brain organoids (unguided type), rather than neurospheres, which can mimic the cytoarchitecture and developmental trajectories found. Importantly, we show that microglia in our CO-iM model can be successfully and preferentially infected with HIV at a particularly high percentage which led to heightened inflammatory responses.

Microglia are known to account for around 5–10% of total cells in the CNS^73^ with astrocytes and neurons being the major cell types at similar proportions, depending on brain region and size of neurons, but typically within 20-40%^74^. In the CO-iM model, on average, around 7.3% of total cells were microglia (double positive for CD11b and Iba1) compared to less than 1% in conventional COs. In addition, astrocytes (around 35-50%) and neurons (around 44-51%) were abundant, and their defining transcripts and antigens were well expressed in CO-iM. Thus, this novel method can generate organoids with physiologically relevant contents of microglia, astrocytes, neurons, and other brain cells providing a platform to study interactions between CNS cell types in a 3D conformation.

CSF1 and IL34 are important proteins for growth and maturation of microglia. Significantly induced expression of *CSF-1R*, a common receptor for both CSF1 and IL34 in combination with strong expression of *IL34* at transcript levels suggests a self-sufficient milieu for survival and maturation of microglia in our CO-iM organoids. Further, significant induction of multiple sensome markers and the general immunity related transcripts by RNAseq in CO-iMs is a compelling indication that microglia are functional in our organoid model. In addition, elevated levels of *C1q* and *C3*, the two main players of initial activation of complement cascade, along with *C3AR1, C7* and *MASP1* suggests that microglia may be playing an important role in synapse pruning and refinement in our CO-iM model. At the same time, heightened complement activation could be detrimental and is associated with cognitive impairment because of excessive pruning^75–77^. CO-iMs exhibited downregulation of some of its components including *C4B* and *C5* suggesting a balanced regulation of pathway to maintain a homeostatic condition. Keeping in mind the high interest of late to develop targeted therapeutics against this pathway to mitigate neurodegeration^78–81^, we believe that CO-iMs could provide a good platform toward high throughput screening of small molecules and drugs that can specifically regulate complement activation. Taken together, we can conclude that microglia in CO-iMs are functional for their known immunity and particular susceptibility to HIV infection.

HIV enters the brain within two weeks of acute infection and within the brain, microglia and macrophages are the major targets of HIV infection. In the post cART era, it is imperative to understand and dissect the role of microglia in HIV neuropathogenesis to gain insights into two major unsolved issues, HAND and HIV latency. Several recent studies have shown that microglia are successfully infected with HIV resulting in abrogated immune responses. Limitations of these studies include either the use of monolayer cultures of microglia^47^, incorporation of externally infected microglia cells into maturing organoids^82^, or use of high amounts of virus particles to infect low abundant microglia that are developed innately in COs^48^. As microglia do not present the well-characterized SAMHD1 restriction to HIV-1 in myeloid cells (macrophages and dendritic cells)^83^, we found them to be infected at a particularly high ratio soon after viral inoculation (within 5 days) where the viruses were added into the culture media and crossed the organoid tissue layers to initiate replication deep into the organoids. We find that when comparing CO-iMs with and without HIV infection, there is a high upregulation of immune markers and their release with infection. We also compared inoculated CO-iMs with and without antiretrovirals that prevent seeding of proviruses into the host genome and found that the transcriptome related to immunity is dependent on viral replication. These allowed us to exclude the effect of extracellular/non-cytoplasmic immune responses such as TLR activation due to the presence of viruses in the inoculum. Interestingly, we find that not all microglia become infected despite the presence of high virus amounts in regions seeded with microglia, denoting a diversity of cells and a possible protective early triggering of innate responses and cytokine milieu from the initial rounds of replication that warrants further investigation. Infection of the CO-iMs result in strong neuroinflammation profiles, both at a transcriptomic (**Figs. 5N and O**) and cytokine level (**Fig. 5M**), and infection results in a transcriptomic profile associated with suppression of normal neural function and development (**Fig. 6**). Despite controlling viral spread and systemic replication, modern antiretroviral therapies do not prevent any of the late-stages of the HIV life cycle as they focus on reverse transcriptase and integrase inhibitors, allowing for the production of nucleic acids, viral protein, and viruses capable of maturation and fusion to other cells^84–87^. Thus, our model provides a unique platform to test specific molecular interactions between HIV particles and the cellular innate immune machinery that detects HIV and is known to mount cascades of immune responses. For example, innate sensing of foreign nucleic acids by cGAS/STING in blood myeloid cells occurs within the first hours of infection followed by a severe down regulation of their activation^88–90^.

The CO-iM model offers a suitable platform where microglia with or without infection can crosstalk with other glial and neuronal cell types in a human-brain-like environment. As the cellular origin of CO-iMs are iPSCs, this is a suitable platform to detail mechanistic functions related to specific genes and to specific cellular lineages (HPCs vs NPCs), as iPSCs are easily genetically manipulated. In the context of HIV, this platform provides a unique opportunity to model the interaction of human microglia with their neuronal environment and renders it suitable for unmasking the role of microglia and its specific genes in viral-related inflammation, its role in HIV persistence under cART, and the biological consequences of infection to astrocytic and neuronal health.

## Author contributions

JIM, SDN, and LA conceptualized and designed the study. SDN, JPZ, JIM performed the study. All interpreted the data. MKA contributed to the generation of organoids, and manuscript editing. AR, TS and SG helped with organoid slicing, qRT-PCR and p24 ELISA, respectively. RLR performed RNAseq and scRNAseq analysis. SDN and JIM wrote the manuscript with input from all authors.

## Acknowledgments

This work was in part supported by grants from the NIH R01DA055497 and R01NS108796 to LA, Walder Foundation innovation Top Up award to JRS, R01MH125778, P01AI169600, and the P30AI117943 Third Coast Center for AIDS Research (TC-CFAR) to RLR, and R21MH129205 and R61DA058348 to JIM. Library preparation and sequencing were performed at Northwestern University NUSeq core. Cytokine panels were performed by the Rush University Biomarker Development core.

