## Supplementary figures and images for "HIV-1 infection promotes neuroinflammation and neuron pathogenesis in novel microglia-containing cerebral organoids"

### Supplemental Figure 1

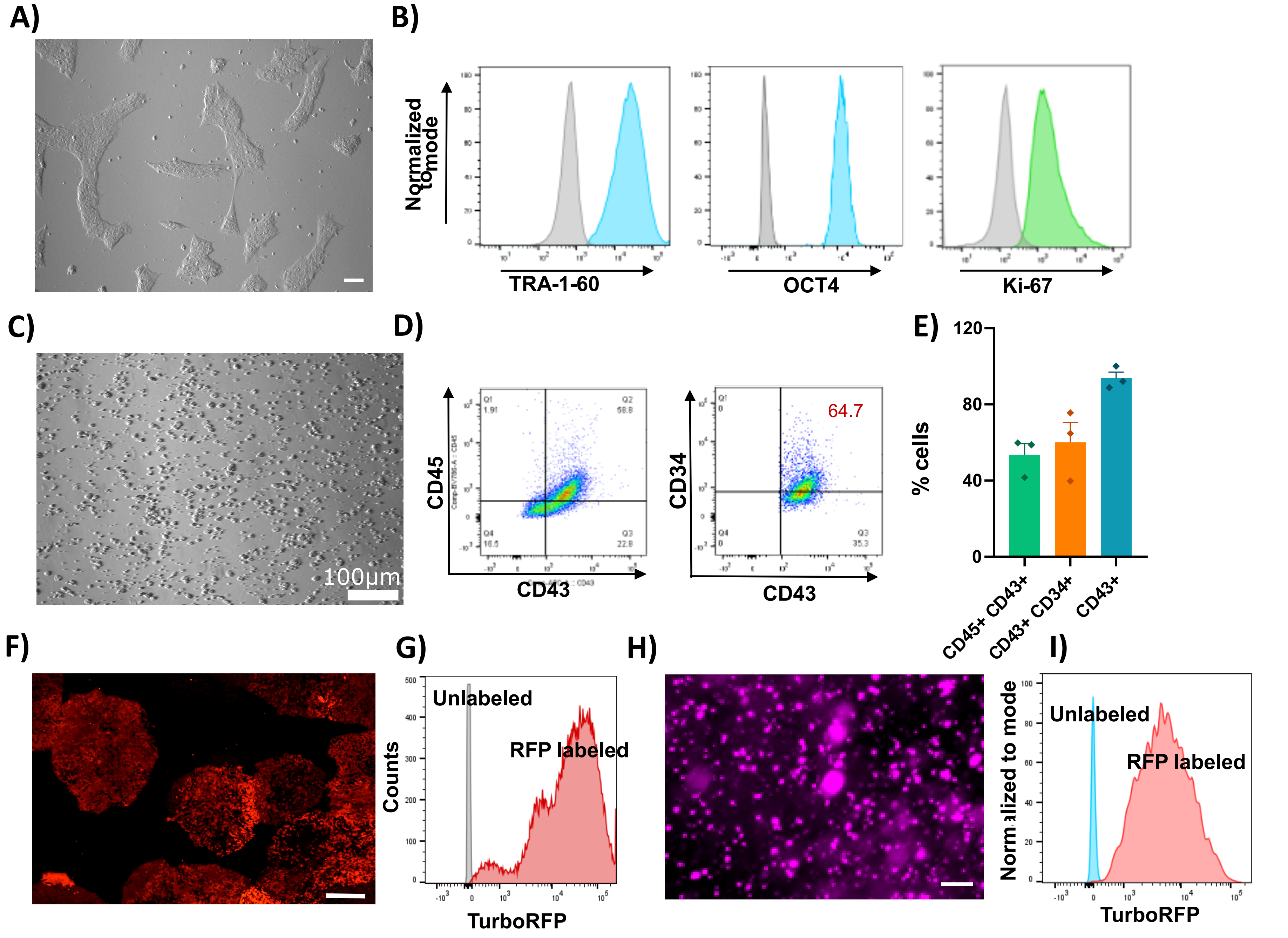

### Supplemental Figure 2

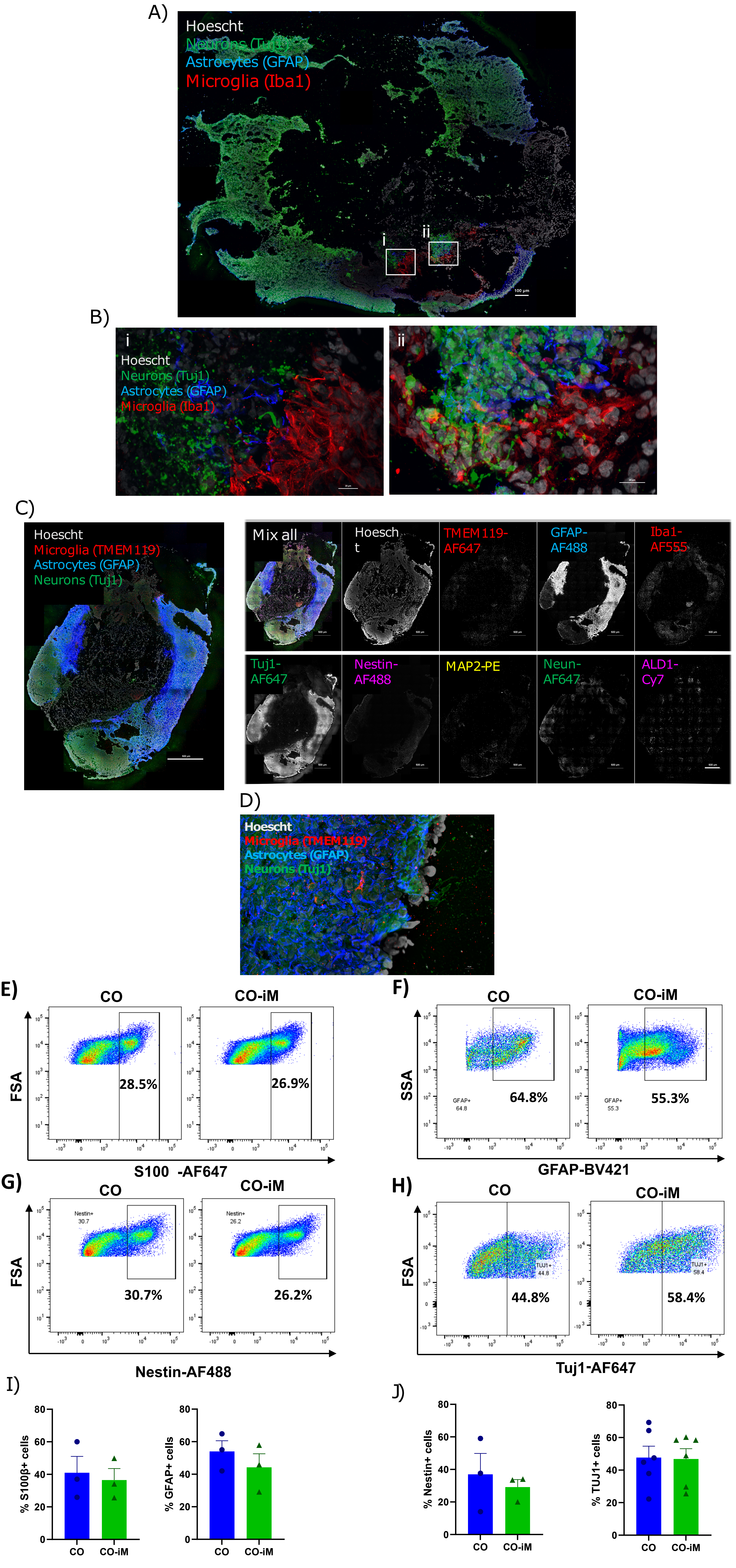

### Supplemental Figure 3

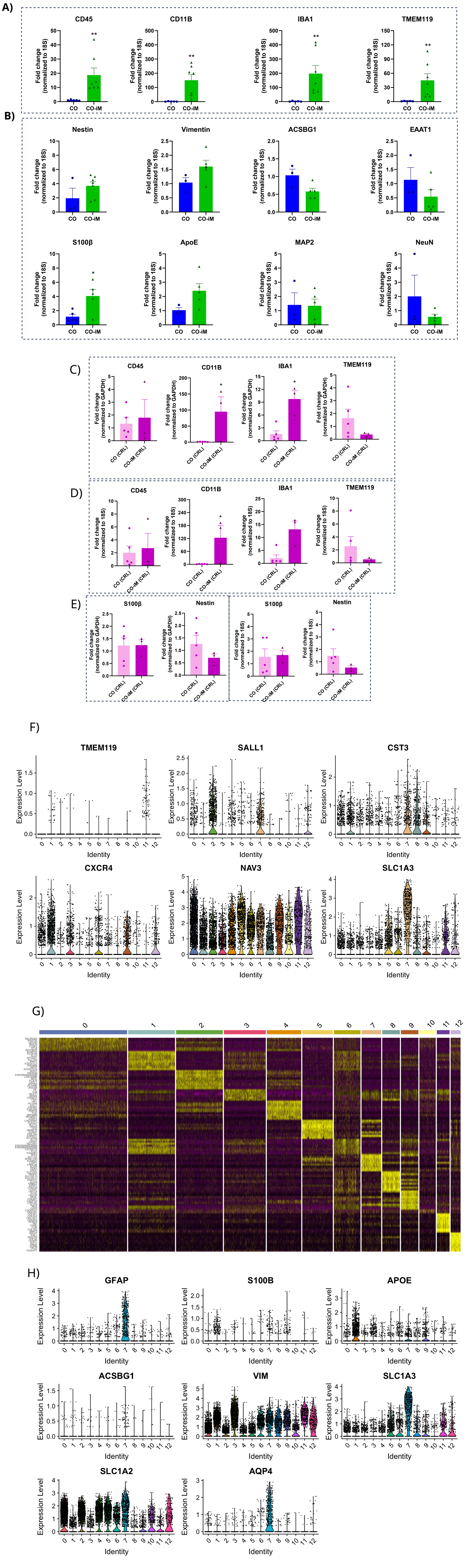

### Supplemental Figure 4

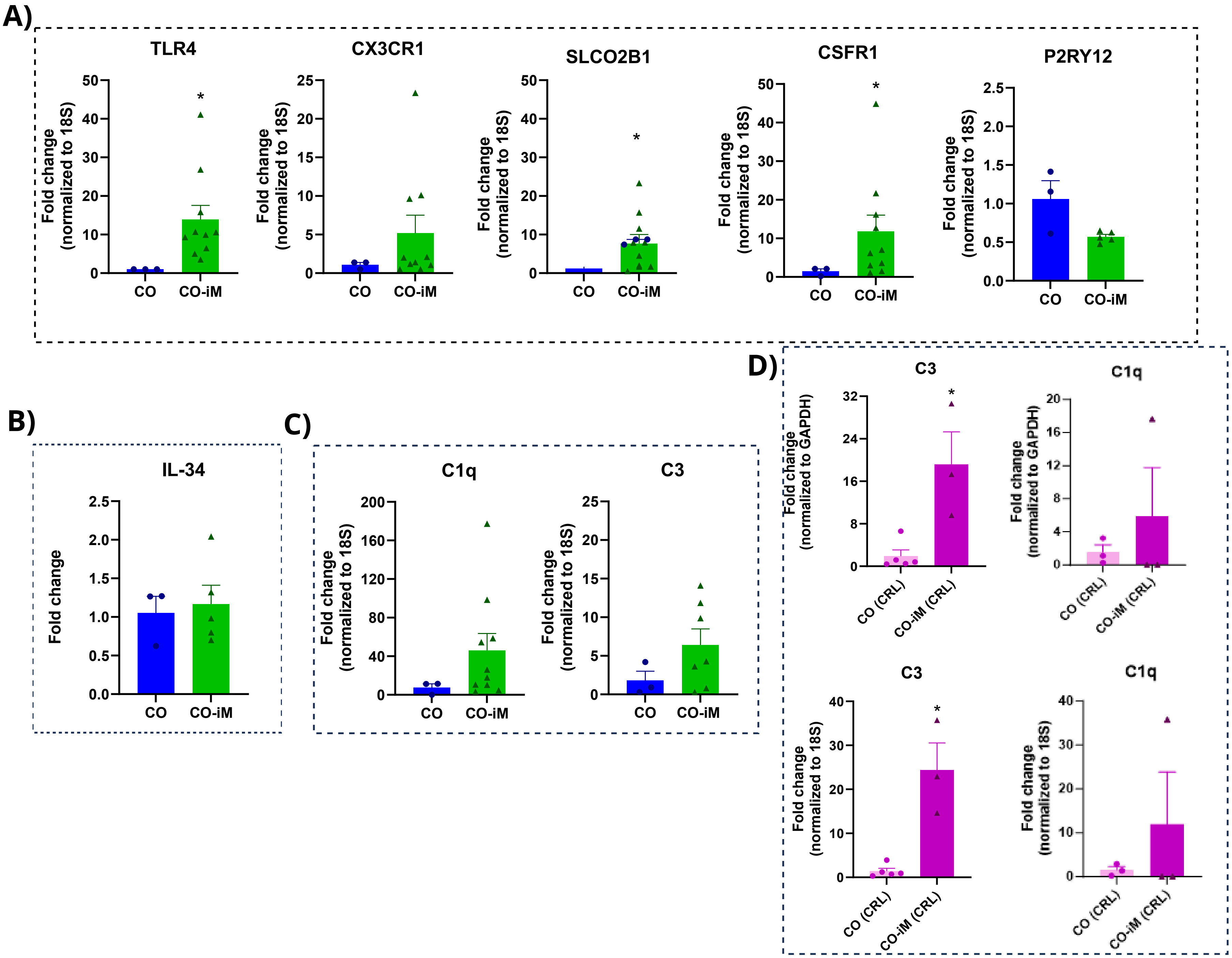

### Supplemental Figure 5

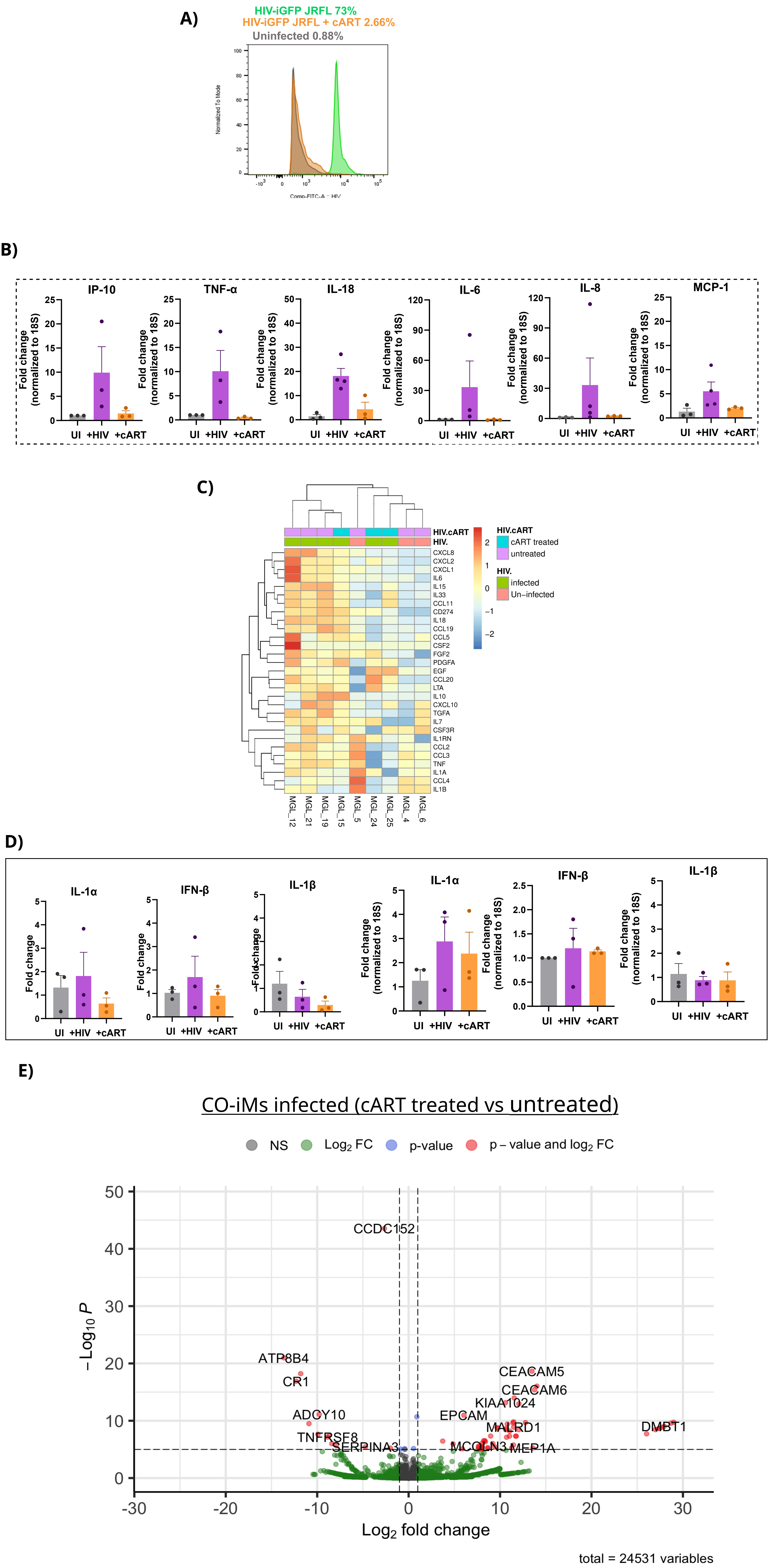
