## Supplemental Figure Legends for "HIV-1 infection promotes neuroinflammation and neuron pathogenesis in novel microglia-containing cerebral organoids"

### Supplementary Figure Legends:

**Supplementary Figure 1: Quality assessment of iPSCs and HPCs. (A)** Representative bright field image of iPSC-Ru. Scale bar, 100 μm. **(B)** Flow cytometry analysis (histogram) indicating iPSCs positive for OCT4, TRA-1-60 (pluripotency markers) and Ki67 (proliferation marker). **(C)** Representative bright field image of HPCs generated from iPSC-Ru. Scale bar, 100 μm. **(D)** Flow cytometry analysis (dot plot) of single cells from (C) indicating positivity for CD45, CD43 and CD34. **(E)** Cumulative data showing percent CD45/CD43, CD43/CD34 double positive as well as CD43 single positive cells in HPCs generated using iPSC-Ru line (n=3). Data are presented as mean ± SD. **(F)** Representative fluorescence image of iPSC-Ru expressing Turbo-RFP stably integrated via lentiviral approach. Scale bar, 100 μm. **(G)** Flow analysis (histogram) of single cells from (F) indicating RFP+ cells. **(H)** Representative fluorescence image of RFP+ HPCs generated from iPSC-Ru stably expressing Turbo-RFP. **(I)** Flow analysis (histogram) of single HPC cells from (H) indicating positivity for RFP.

**Supplementary Figure 2: Phenotypic characterization informing enrichment of microglia in CO-iMs. (A)** Representative mIF image of a CO-iM stained for nucleus (Hoescht, white), microglia (Iba1, red), neurons (TUJ1, green) and astrocytes (GFAP, blue) Scale bar, 100 μm. **(B)** Zoomed in insets of i), and ii) regions Scale bar, 100 μm. **(C)** Left, representative mIF image of an another CO-iM stained for nucleus (Hoescht, white), microglia (TMEM119, red), neurons (TUJ1, green) and astrocytes (GFAP, blue) Scale bar, 100 μm. Right, panel showing individual staining for indicated markers and a merged image from all. Scale bar, 100 μm. **(D)** mIF image of a CO-iM showing microglia (TMEM119, red), astrocytes (GFAP, blue) and neurons (TUJ1, green). Scale bar, 100 μm. **(E-H)** Representative dot plots showing S100β, GFAP, Nestin and Tuj1 positive populations from total cells in CO and CO-iMs, respectively. **(I and J)** Cumulative data indicating percent S100β, GFAP, Nestin and Tuj1 positive cells in CO and CO-iMs generated using iPSC-Ru line (n=3-6). Each symbol represents an individual organoid. Data are presented as mean ± SD using non-parametric Mann-Whitney test. *p < 0.05, **p < 0.01, ***p < 0.001.

**Supplementary Figure 3: Genotypic characterization revealing enrichment of microglia markers in CO-iMs**. qRT-PCR analysis of COs (n=3-5) and CO-iMs (n=5-7). mRNA levels were normalized to the endogenous reference *18S* and expressed as fold change relative to CO using $2^{-(\Delta\Delta Ct)}$. Fold change for each sample in COs was assessed by subtracting its ΔCt from average ΔCt, followed by $2^{-(\Delta\Delta Ct)}$. **(A)** mRNA levels of homeostatic microglia markers: *CD45*, *CD11B*, *IBA1* and *TMEM119*. **(B)** mRNA levels of NPC, astrocyte and mature neuronal markers as indicated in the text. **(C and D)** mRNA levels of *CD45*, *CD11B*, *IBA1* and *TMEM119* normalized to *GAPDH* or *18S* in COs (n=5) and CO-iMs (n=3) generated using iPSC-CRL. **(E)** mRNA levels of *S100β* and *Nestin* normalized to *GAPDH* or *18S* in COs (n=5) and CO-iMs (n=3) generated using iPSC-CRL.

Each symbol represents an individual organoid. Data are presented as mean ± SD using non-parametric Mann-Whitney test. *p < 0.05, **p < 0.01, ***p < 0.001. (**F**) Violin and dotplots showing single cell relative expression levels for microglia representative transcript markers for each of the Seurat clusters displayed in figure 4. (**G**) Top 10 hits per Seurat clusters displayed in Figure 4. (**H**) Violin and dotplots showing single cell relative expression for astrocyte representative transcript markers for each of the Seurat cluster displayed in figure 4.

**Supplementary Figure 4: Microglia sensome and complement cascade pathway characterization in CO-iMs at mRNA level.** qRT-PCR analysis of COs and CO-iMs to assess mRNA levels computed using $2^{-(\Delta\Delta Ct)}$. Fold change for each sample in COs was assessed by subtracting its delta Ct from average delta Ct, followed by $2^{-(\Delta\Delta Ct)}$. **(A)** mRNA levels of microglia immune sensome markers: *CX3CR1*, *SLCO2B1*, *TLR4*, *CSF1-R* and *P2RY12.* **(B)** mRNA levels of *IL34,* an essential cytokine for microglia maturation and survival and **(C)** mRNA levels of *C3* and *C1Q*, components of complement pathway in CO and CO-iMs generated using iPSC-Ru. Ct values normalized to 18S. (n=3 for COs and n=5-7 for CO-iMs). **(D)** mRNA levels of *C3* and *C1Q*, in CO and CO-iMs generated using iPSC-CRL. Ct values normalized to 18S or GAPDH. (n=5 for COs and n=3 for CO-iMs).

Each symbol represents an individual organoid. Data are presented as mean ± SD using non-parametric Mann-Whitney test. *p < 0.05, **p < 0.01, ***p < 0.001.

**Supplementary Figure 5: Robust inflammatory response of CO-iMs to HIV-1 infection. (A)** CO-iMs were infected with HIV-iGFP_JRFL, mock infected or infected and treated with cART for 5 days as explained in the methods section, dissociated into single cell suspension and stained for cell surface microglia markers, (CD45 and CD11b) and HIV (GFP) and analyzed via flow cytometry and plotted as a histogram. n=1 for each group (mock, infected and infected+cART). **(B)** qPCR analysis for mRNA expression levels of various pro-inflammatory cytokines/chemokines (as indicated in the figure) in CO-iMs either infected with $HIV_{BaL}$ for 5 days, or mock infected, or infected and treated with cART. n=3 in each category. **(C)** A heatmap showing expression levels of a number of pro-inflammatory cytokines/chemokines in mock vs infected vs infected+cART treated CO-iMs, (n = 3 for each condition). Color indicates the expression level normalized for each gene using a Z-score. **(D)** qPCR analysis for mRNA expression levels of various pro-inflammatory cytokines/chemokines (as indicated in the figure) in CO-iMs either infected with $HIV_{BaL}$ for 5 days, or mock infected, or infected and treated with cART. n=3 in each category. **(E)** Volcano plot comparison of infected vs infected+cART treated CO-iMs indicating average log2 (fold change) versus log10 (FDR) for all genes. Genes upregulated and downregulated by 2-fold change and FDR < 0.05 are labeled with red dots.

Each symbol represents an individual organoid. Data are presented as mean ± SD using non-parametric Mann-Whitney test. *p < 0.05, **p < 0.01, ***p < 0.001.
