## Supplementary material for "HIV-1 infection promotes neuroinflammation and neuron pathogenesis in novel microglia-containing cerebral organoids": Tables S1 S2 S3

**Table S1. List of antibodies used for flow cytometry in this study.**

| Antigen | Clone | Type | Fluorochrome | Identifier | Source |
|---|---|---|---|---|---|
| TRA-1-60 | TRA-1-60R | Mouse monoclonal | PE | 60064PE.1 | Stem Cell Technologies |
| OCT4(OCT3) | 3A2A20 | Mouse monoclonal | PE | 60093PE.1 | Stem Cell Technologies |
| OCT4(OCT3) | 3A2A20 | Mouse monoclonal | AF488 | 60093AD.1 | Stem Cell Technologies |
| CD43 | CD43-10G7 | Mouse monoclonal | APC | 343206 | BioLegend |
| CD34 | 581 | Mouse monoclonal | PE | 343506 | BioLegend |
| CD45 | HI30 | Mouse monoclonal | BV786 | 304048 | BioLegend |
| Iba1 | EPR6136(2) | Rabbit monoclonal | PE | ab209942 | Abcam |
| CD11b | M1/70 | Mouse monoclonal | AF700 | 557918 | BD Biosciences |
| S100β | C-3 | Mouse monoclonal | AF647 | sc-393919 | Santa Cruz |
| GFAP | 1B4 | Mouse monoclonal | BV421 | 560297 | BD Biosciences |
| Tubulin β3 | TUJ1 | Mouse monoclonal | AF647 | 801210 | BioLegend |
| Nestin | 10C2 | Mouse monoclonal | AF488 | 60091AD.1 | Stem Cell Technologies |
| HIV-1-Gag | KC57 | Mouse monoclonal | FITC | 6604665 | Beckman Coulter |

**Table S2. List of primers used for qRT-PCR in this study.**

| Gene | Forward Primer | Reverse Primer |
|---|---|---|
| *GAPDH* | 5'-ACC-ACC-CTG-TTG-CTG-TAG-CCA-AAT | 5'-TGA-CTT-CAA-CAG-CGA-CAC-CCA-CT |
| *18S* | 5'-GTA-ACC-CGT-TGA-ACC-CCA-TT | 5'-CCA-TCC-AAT-CGG-TAG-TAG-CG |
| *CD45* | 5'-CTG-GAG-GAT-GAT-TTG-GGA-ACA | 5'-CTT-CCA-TTG-ACG-GCC-AGT-AT |
| *CD11B* | 5'-GGA-ACG-CCA-TTG-TCT-GCT-TTC-G | 5'-ATG-CTG-AGG-TCA-TCC-TGG-CAG-A |
| *IBA1* | 5'-CCC-TCC-AAA-CTG-GAA-GGC-TTC-A | 5'-CTT-TAG-CTC-TAG-GTG-AGT-CTT-GG |
| *TMEM119* | 5'-TCC-AGG- GTC-AGA-TTA-CAA-GAG-CAC | 5'-ACT-GTT-GAT-TCT-GGA-GGG-TTT-GA |
| *S100β* | 5'-GAA-GAA-ATC-CGA-ACTG-AAG-GAG-C | 5'-TCC-TGG-AAG-TCA-CAT-TCG-CCG-T |
| *ACSBG1* | 5'-CCC-CTT-GAC-CTG-TGA-TGA-CC | 5'-GAG-ACG-GGA-TGG-ACT-TGG-A |
| *EAAT1* | 5'-GGT-TGC-TGC-AAG-CAC-TCA-TCA-C | 5'-CAC-GCC-ATT-GTT-CTC-TTC-CAG-G |
| *Vimentin* | 5'-GAT-TCA-CTC-CCT-CTG-GTT-GAT-AC | 5'-GTC-ATC-GTG-ATG-CTG-AGA-AGT |
| *Nestin* | 5'-TCA-AGA-TGT-CCC-TCA-GCC-TGG-A | 5'-AAG-CTG-AGG-GAA-GTC-TTG-GAG-C |
| *MAP2* | 5'-CAG-GAG-ACA-GAG-ATG-AGA-ATT-CC | 5'-CAG-GAG-TGA-TGG-CAG-TAG-AC |
| *NEUN* | 5'-CCG-AGT-GAT-GAC-CAA-CAA-GAA | 5'-CAT-AGA-ATT-CAG-GCC-CGT-AGA-C |
| *APOE* | 5'- GGG-TCG-CTT-TTG-GGA-TTA-CCT-G | 5'- CAA-CTC-CTT-CAT-GGT-CTC-GTC-C |
| *CX3CR1* | 5'-CAC-AAA-GGA-GCA-GGC-ATG-GAA-G | 5'-CAG-GTT-CTC-TGT-AGA-CAC-AAG-GC |
| *SLCO2B1* | 5'-GTT-TCG-GCG-AAA-GGT-CTT-CTT-CGC-AG | 5'-CCA-TCC-TGC-TTC-TTC-GTG-GAC-T |
| *TLR4* | 5'-CCC-TGA-GGC-ATT-TAG-GCA-GCT-A | 5'-AGG-TAG-AGA-GGT-GGC-TTA-GGC-T |
| *CSFR1* | 5'-GCT-GCC-TTA-CAA-CGA-GAA-GTG-G | 5'-CAT-CCT-CCT-TGC-CCA-GAC-CAA-A |
| *P2RY12* | 5'-GTG-TCA-AGT-TAC-CTC-CGT-CAT-A | 5'-TAA-ATG-GCC-TGG-TGG-TCT-TC |
| *IL-34* | 5'-CCA-AGG-TGG-AAT-CCG-TGT-TGT-C | 5'-CAC-CTC-ACA-GTC-CTG-CCA-GTT-T |
| *C3* | 5'-GAC-ATT-CCG-GAA-CTC-GTC-AA | 5'-CGT-ACT-CCT-TCA-CCT-CAA-ACT-C |
| *C1Q* | 5'-CAA-CAC-AGG-CTG-CTA-CGG-GAT-C | 5'-CTG-CCC-TTT-GGG-TCC-TCG-GAT |
| *IP-10* | 5'-GGT-GAG-AAG-AGA-TGT-CTG-AAT-CC | 5'-GTC-CAT-CCT-TGG-AAG-CAC-TGC-A |
| *TNFα* | 5'- CTCTTCTGCCTGCTGCACTTTG | 5'- ATGGGCTACAGGCTTGTCACTC |
| *IL-18* | 5'-GAT-AGC-CAG-CCT-AGA-GGT-ATG-G | 5'-CCT-TGA-TGT-TAT-CAG-GAG-GAT-TCA |
| *1L-6* | 5'-AGA-CAG-CCA-CTC-ACC-TCT-TCA-G | 5'-TTC-TGC-CAG-TGC-CTC-TTT-GCT-G |
| *IL-8* | 5'-GAG-AGT-GAT-TGA-GAG-TGG-ACC-AC | 5'-CAC-AAC-CCT-CTG-CAC-CCA-GTT-T |
| *MCP-1* | 5'-AGA-ATC-ACC-AGC-AGC-AAG-TGT-CC | 5'-TCC-TGA-ACC-CAC-TTC-TGC-TTG-G |
| *IL-1α* | 5'-TGT-ATG-TGA-CTG-CCC-AAG-ATG-AAG | 5'-AGA-GGA-GGT-TGG-TCT-CAC-TAC-C |
| *INF-β* | 5'- CTT-GGA-TTC-CTA-CAA-AGA-AGC-AGC | 5'- TCCTCCTTCTGGAACTGCTGCA |
| *IL-1β* | 5'-CCA-CAG-ACC-TTC-CAG-GAG-AAT-G | 5'-GTG-CAG-TTC-AGT-GAT-CGT-ACA-GG |

**Table S3. List of antibodies used for immunofluorescence in this study.**

| Antigen | Clone | Type | Fluorochrome | Identifier | Source |
|---|---|---|---|---|---|
| HIV-1 Gag | KC57 | Mouse monoclonal | PE | 6604667 | Beckman Coulter |
| HIV-1 CA | 241D | Human monoclonal | Unconjugated | ARP-1244 | HIV reagents |
| HIV-1 CA | 71-31 | Human monoclonal | Unconjugated | ARP-530 | HIV reagents |
| Iba1 | E404W | Rabbit monoclonal | AF555 | 36618 | Cell Signaling |
| Iba1 | EPR6136(2) | Rabbit monoclonal | PE | ab209942 | Abcam |
| CD11b | ICRF44 | Rabbit monoclonal | APC-Cy7 | 557754 | BD Biosciences |
| TMEM119 | 106-6 | Mouse monoclonal | AF647 | Ab225494 | Abcam |
| S100β | C-3 | Mouse monoclonal | AF488 | sc-393919 | Santa Cruz |
| GFAP | 1B4 | Mouse monoclonal | AF488 | 560297 | BD Biosciences |
| GFAP | GA5 | Mouse monoclonal | AF647 | 3657S | Cell Signaling |
| Tubulin b3 | TUJ1 | Mouse monoclonal | AF647 | 801210 | BioLegend |
| Nestin | 10C2 | Mouse monoclonal | AF488 | 60091AD.1 | Stem Cell Technologies |
| NeuN | EPR12763 | Rabbit monoclonal | AF647 | Abcam | Abcam |
| MAP2 | AP20 | Mouse monoclonal | PE | sc-32791 | Santa Cruz |
| ALDH1L1 | 2E7 | Mouse monoclonal | NHS-ester-Cy7 | MA5-47403 | Stem Cell Technologies |
